# Stacking haplotypes from the Vavilov wheat collection to accelerate breeding for multiple disease resistance

**DOI:** 10.1101/2024.03.28.587294

**Authors:** Jingyang Tong, Zerihun T. Tarekegn, Samir Alahmad, Lee T. Hickey, Sambasivam K. Periyannan, Eric Dinglasan, Ben J. Hayes

## Abstract

Wheat production is threatened by numerous fungal diseases, but the potential to breed for multiple disease resistance (MDR) mechanisms is yet to be explored. Here, significant global genetic correlations and underlying local genomic regions were identified in the Vavilov wheat diversity panel for six major fungal diseases, including biotrophic leaf rust (LR), yellow rust (YR), stem rust (SR), hemibiotrophic crown rot (CR), and necrotrophic tan spot (TS) and Septoria nodorum blotch (SNB). By adopting haplotype-based local genomic estimated breeding values, derived from an integrated set of 34,899 SNP and DArT markers, we established a novel haplotype catalogue for resistance to the six diseases in over 20 field experiments across Australia and Ethiopia. Haploblocks with high variances of haplotype effects in all environments were identified for three rusts and pleiotropic haploblocks were identified for at least two diseases, with four haploblocks affecting all six diseases. Through simulation we demonstrated that stacking optimal haplotypes for one disease could improve resistance substantially, but indirectly affected resistance for other five diseases, which varied depending on the genetic correlation with the non-target disease trait. On the other hand, our simulation results combining beneficial haplotypes for all diseases increased resistance to LR, YR, SR, CR, TS and SNB, by up to 48.1%, 35.2%, 29.1%, 12.8%, 18.8% and 32.8%, respectively. Overall, our results highlight the genetic potential to improve MDR in wheat. The haploblock-based catalogue with novel forms of resistance provides a useful resource to guide desirable haplotype stacking for breeding future wheat cultivars with MDR.

## Introduction

Wheat (*Triticum aestivum*) production is constantly threatened by numerous biotrophic and necrotrophic fungal diseases (Hafeez et al., 2021). Major fungal diseases include leaf rust (LR, caused by *Puccinia triticina* f. sp. *tritici*), stripe rust (YR, caused by *P. striiformis* f. sp. *tritici*), stem rust (SR, caused by *P. graminis* f. sp. *tritici*), powdery mildew (caused by *Blumeria graminis* f. sp. *tritici*), tan spot (TS, caused by *Pyrenophora tritici*-*repentis*), Septoria nodorum blotch (SNB, caused by *Parastagonospora nodorum*), Fusarium head blight (mainly caused by *Fusarium graminearum*) and Fusarium crown rot (CR, mainly caused by *Fusarium pseudograminearum*), which cause severe yield and economic losses globally (Figueroa et al., 2018; Savary et al., 2019). To manage these diseases, the deployment of host plant-mediated genetic resistance is preferred over chemical control due to its cost effectiveness, reliability, and environmentally friendly nature (Jørgensen et al., 2017). Breeding of resistant wheat cultivars has been accelerated by a multitude of identified genetic loci or genes providing resistance to individual diseases, as well as in-depth understanding of pathogen-plant interactions and resistance/susceptibility mechanisms (Figueroa et al., 2018; Liu et al., 2020; Pal et al., 2022; Peters Haugrud et al., 2022; Su et al., 2021). Despite significant progress to clone genes and breeding success for resistance to individual wheat diseases, insights are still limited into the nature of the genetic relationships between multiple disease resistance traits that might have correlated genomic regions (Liu et al., 2020; Zhang et al., 2021). Some examples demonstrate pleiotropic resistance, such as *Lr34/Yr18/Sr57/Pm38/Ltn1* (Krattinger et al., 2009), *Lr46/Yr29/Sr58/Pm39/Ltn2* (Singh et al., 1998), *Lr67/Yr46/Sr55/Pm46/Ltn3* (Moore et al., 2015) and *Sr2/Lr27/Yr30/Pbc1* (Mago et al., 2011) with resistance to more than three diseases. To support wheat breeding for multiple disease resistance (MDR), genetic studies examining resistance to multiple diseases, each caused by distinct pathogen is important (Nene, 1988) and could inspire efficient strategies for combining multiple and complex traits in crop breeding (Grotzinger et al., 2019; Song et al., 2023).

The deployment of a single major resistance (*R*) gene can provide high levels of resistance to a specific disease, however prone to breaking down due to intense selection pressure on the pathogen to evolve strains with mutation in the corresponding virulence gene, required for resistance (Sánchez-Martín and Keller, 2021). Quantitative disease resistance (QDR) relying on multiple additive major and minor *R* genes and their effective combinations is widely recognised as a more durable host defense mechanism (Singh et al., 2014). For example, strong background resistance resulting from the accumulation of multiple minor genes has been critical to success in rust resistance breeding efforts at CIMMYT (Bhavani et al., 2019). These minor genes can also enhance and prolong the efficiency of qualitative resistance, suggesting that stacking both minor and major *R* genes simultaneously could be beneficial (Brun et al., 2010). Extending the gene pyramiding concept of Hospital and Dekkers (1994), several strategies have been proposed to identify and stack chromosome segments defined by marker-based haplotypes with favourable estimated genetic effects on target traits (Goiffon et al., 2017; Kemper et al., 2012; Voss-Fels et al., 2019). Employing a haplotype-based strategy could enhance the efficiency of developing durable resistance and MDR in a wheat breeding programme since it enables the tracking and selection of blocks of markers in linkage disequilibrium (LD), rather than a single marker which is more prone to breaking down in association with the trait through recombination (Voss-Fels et al., 2019). However, the first step is to discover favourable haplotypes with strong effects on resistance to a range of diseases (Wiesner-Hanks and Nelson, 2016).

A historically important wheat germplasm collection is preserved at the N.I. Vavilov Institute of plant genetic resources (VIR), which was collected by Russian botanist N.I. Vavilov and his colleagues in the early 1900s (Mitrofanova, 2012). Through our previous studies, we showed the Vavilov wheat diversity panel provided a useful source of genetic resistance to diseases, such as LR (Riaz et al., 2018), YR (Jambuthenne et al., 2022), TS (Dinglasan et al., 2019) and SNB (Phan et al., 2018). However, these studies used a single marker genome-wide association study (GWAS)-approach and examined resistance to each disease in isolation. Thus, the implications for haplotype-based breeding and its potential to breed for MDR is yet to be explored.

The aim of this study was to apply a haplotype-based approach to the Vavilov wheat diversity panel to: (1) explore global and local (haploblock) genetic correlations between resistance for six wheat major diseases, i.e. LR, YR, SR, TS, SNB, and CR; (2) establish a haplotype resistance catalogue for each respective disease using haploblocks across the wheat genome, and to discover key MDR haploblocks; (3) assess the genetic potential of MDR and explore the most desirable pathway to maximise MDR via haplotype stacking.

## Results

### Vavilov wheat accessions are genetically diverse with rapid LD decay

A panel of 295 Vavilov wheat accessions with passport information (Table S1) was genotyped with three different marker platforms, resulting in 34,899 high-quality markers. This included 11,992 (34.36%) from a 90K SNP array, 3,325 (9.53%) from DArT SNP, and 19,582 (56.11%) from DArTsilico markers. These markers were used to construct an integrated marker map with reliable physical positions using the IWGSC RefSeq v2.1 (Figure S1 & S2). The total length of the marker map was 14,216 Mb, and average marker density, missing rate and minor allele frequency was 0.41 Mb/marker, 0.10, and 0.28, respectively, suggesting high-density and good marker quality following marker integration (Table S2). The B subgenome had the longest length and highest marker density, followed by the A and D subgenomes.

Population structure and diversity were investigated by the genotypic data and different approaches (Figure S3). According to the hierarchical cluster, genetic similarity and the matrices corresponding to two to four hypothetical subgroups derived from the STRUCTURE analysis, three subpopulations were inferred to be the best grouping classified by the dashed horizontal lines in Figure S3. The results were somewhat different from Riaz et al., (2017), which grouped the Vavilov panel into two major subpopulations mainly based on wheat cultivation status, probably due to different genotypic data (Riaz et al., 2017, only used DArTsilico markers) and the clustering methods. Agreeing with Figure S3, three subpopulations were clearly classified based on the first three principal components, as well as in the two-dimension principal component analysis (PCA) and Neighbor-Joining (NJ) tree analysis (Figure S4). Subpopulation 1 included 54 Asian wheat landraces with spring growth habit, mainly originated from India and Pakistan; subpopulation 2 consisted of 100 accessions included landraces and cultivars mainly from Asia (e.g. Pakistan, China, Armenia, Kazakhstan); subpopulation 3 comprising 141 accessions was mainly represented by landraces and cultivars from Europe (e.g. Russia, Ukraine) (Table S1). Given that understanding LD is important in haplotype-based strategy, we estimated average LD decay distances in the entire panel. Results showed that whole-genome average LD decay distance was 1.25 Mb, while A, B, D subgenomes were 1.22, 0.98 and 1.37 Mb, respectively, implying a rapid LD decay (Figure S5).

Taken together, we demonstrate high genetic diversity and rapid LD decay in the Vavilov wheat panel.

### Screening against multiple diseases revealed association between resistance of six wheat major diseases

A total of 22 phenotypic datasets for resistance to biotrophic (LR, YR, SR), necrotrophic (TS, SNB), and hemibiotrophic (CR) fungal diseases produced from various field environments (except SNB) in Queensland (QLD) and Ethiopia were used in this study (Table S3). To ensure that all haplotype effects were comparable across multiple diseases, the phenotypic data from different experiments were converted into a consistent scale 0–9 (0 = most resistant; 9 = most susceptible) to enable further analyses (Figure S6). A wide range of almost continuous distributions of resistance were noticed in the Vavilov diversity panel for different diseases across multiple environments (Figure S6). To effectively simplify or reduce the dimensions of the many phenotypic datasets, we tried three different dimension reduction approaches, including relative disease index (Ri), best linear unbiased estimate (BLUE), and eigenvector principal component 1 (PC1). Eigenvector PC1 explained considerable phenotypic variance ranging from 63.6% to 76.1% across multiple field environments (Figure S7). Ri, PC1 and BLUE values describing the resistance level for each disease of each wheat accession have different levels of variation (Figure S8).

Pearson correlations and global genetic correlations were calculated between the six disease resistance traits using the three parameters, respectively (Figure S9). Genetic correlations along the BLUE values were used to represent the genetic relationships between traits. Interestingly, the resistance traits of the three rust diseases showed positive and moderate genetic correlations (*r* = 0.18–0.34) with each other (Figure S9b; Figure 1a). Positive genetic correlation was also observed between TS and YR (*r* = 0.22), as well as CR and SR (*r* = 0.20) resistances. However, resistance to SNB was found to be genetically negatively correlated with LR (*r* = −0.31) and YR (*r* = −0.21).

**Figure 1.**
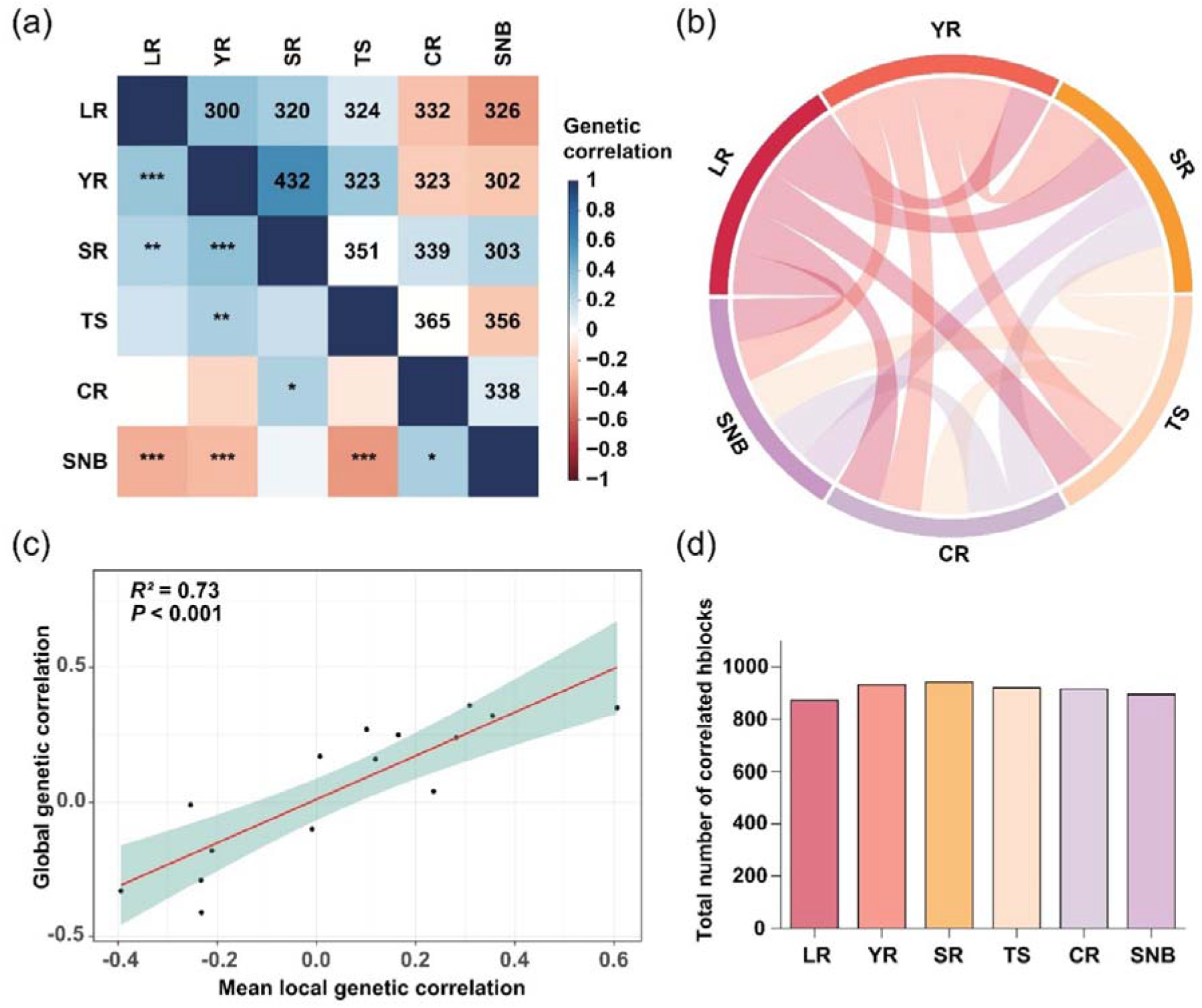
Genetic correlations between the six wheat major diseases, including leaf rust (LR), yellow rust (YR), stem rust (SR), tan spot (TS), crown rot (CR) and Septoria nodorum blotch (SNB). **(a)** Comparison between local genetic correlations (top) and global genetic correlations (bottom). The numbers denote significant correlated haploblocks. The asterisks indicate a significant global genetic correlation. * *P* < 0.1, ** *P* < 0.01, *** *P* < 0.001. **(b)** Chord diagram illustrating the number of significant correlated haploblocks between all trait pairs. The thickness of each chord denotes the number. **(c)** Relationship between the mean correlations of significant correlated haploblocks and the global genetic correlations in all trait pairs. **(d)** Bar plot showing the total number of significant correlated haploblocks detected per disease trait.

In each environment, Pearson correlation analysis and PCA were performed on resistance responses to LR, YR, SR, TS, SNB, and CR (Figure S10). For each disease, the correlation coefficient between different environments was very high (Figure S11). Resistance to SR_21DZ was negatively correlated with LR_14/16QLD, but positively correlated with LR_15QLD. Response against SNB was negatively correlated with LR and YR in QLD but showed little relationship with resistance to rust recorded in Ethiopia. Interestingly, YR showed high correlations within Ethiopia and QLD environments, respectively, but weak to moderate correlations observed between Ethiopia and QLD for YR.

Overall, both phenotypic and genetic relationships were clearly evident and varied among the resistance response against different fungal diseases in the Vavilov wheat panel.

### Local (haploblock-based) genetic correlations between disease resistance traits

Considering the overall genetic relationships between disease resistance traits, we hypothesised that there may be local genomic regions underpinning the global genetic correlations, which can either be in the same direction as the global correlation estimate (‘drivers’), or less commonly in the opposite direction (‘antagonizers’) (Olasege et al., 2022). To explore local genetic correlations, we adopted a novel haploblock-based approach that correlated genetic effects of markers in strong LD. In total, 19,725 genome-wide haploblocks (criteria for grouping adjacent markers into a haploblock was *r*^2^ >= 0.5) were constructed from 34,899 markers in the Vavilov wheat diversity panel and distribution patterns of the defined blocks were displayed across all the 21 wheat chromosomes (Figure S12; Table S4). The number of haploblocks, average number of markers per block and average block size varied between regions, chromosomes and subgenomes (Figure S13). The genome-wide 1,756 haploblocks each harbouring >3 markers were used to calculate local correlations between all 15 pairs of disease resistance traits (Figure S14; Table S5). The results from a sliding-window-based approach, correlation scan (Olasege et al., 2022), was used to compare with that of the haploblock-based method, and similar estimates of local genetic correlations for three resistance trait pairs were observed on most chromosomes between the two methods (Figure S15).

Hundreds of significantly correlated haploblocks were identified in all 15 resistance trait pairs that were either significantly correlated or not correlated in the global analysis (Figure 1a; Figure 1b). However, regression analysis showed that the global genetic correlations were highly concordant (*R*^2^ = 0.73) with mean correlations of significant haploblocks in all trait pairs (Figure 1c), suggesting that higher global genetic correlations were associated with higher mean local genetic correlations. The total number of significantly correlated haploblocks detected per disease resistance trait was similar (Figure 1d), implying that considerable numbers of haploblocks had correlated genetic effects on multiple pairs of disease resistance traits. We investigated three types of haploblocks: ‘driver’ blocks, ‘antagonizing’ blocks and ‘neutral’ blocks that significantly drive, antagonize or do not influence the global genetic correlation, respectively (Figure S16a). Interestingly, the absolute value of global genetic correlation between each trait pair was proportional to the number of ‘driver’ haploblocks (Figure S16b).

In conclusion, we identified local genomic regions that underpinned observed overall genetic correlations among the resistance to different diseases and revealed that a high frequency of ‘driver’ haploblocks were detected for trait pairs with higher global genetic correlations.

### A genome-wide haplotype catalogue with novel forms of resistance discovers genomic regions associated with multi-disease resistance

We used a haplotype-based mapping approach to calculate local GEBVs and variances of haplotypes to identify the chromosomal segments with moderate to large effects on disease resistance. A haplotype resistance catalogue was established with detailed information on genome-wide haplotype effects and corresponding haploblock variances for resistance to the six different diseases across 22 environments (Table S6). Using this resistance catalogue, the top 100 haploblocks (top 0.5%) with the highest variance were identified for each trait in each environment. To identify haploblocks explaining genetic variance across all environments, blocks with consistently large variances (termed “consistent” blocks from herein) that were reported in all environments were identified for each rust (i.e. 3 environments for LR, 11 for YR, and 4 for SR). This revealed 22, 6 and 24 consistent blocks for LR, YR, and SR resistance, respectively (Table S7-S9). Additionally, we used the resistance catalogue to investigate haploblocks accounting for high correlations for YR within and between QLD and Ethiopia environments (Figure S11; Figure S17a). Interestingly, quite a few consistent haploblocks were detected contributing to correlated YR responses within QLD and within Ethiopia environments (Figure S17b). Of these, a subset of 40 haploblocks were classified as consistent blocks in both QLD and Ethiopia (Figure S17b; Table S10).

To explore genetic regions associated with multi-disease resistance, genome-wide haploblock variances were calculated for each resistance trait (Table S6; Figure S18). By using the 100 haploblocks with the highest variances for each respective disease, a total of 158 pleiotropic haploblocks were identified to be potentially relevant to resistance for at least two diseases (Table S11). Upset and Venn diagrams were generated to show the number of overlaps with pleiotropic resistance (Figure 2a). Notably, four haploblocks were identified with high variances for resistance to all six wheat diseases. Meanwhile, the chromosome distributions of the top 100 haploblocks for each disease were displayed on all three subgenomes (Figure 2b & S19). Interestingly, the four haploblocks related to six diseases were all located within the A subgenome, particularly 2A chromosome, indicating this subgenome as an important source of MDR in wheat (Figure 2b).

**Figure 2.**
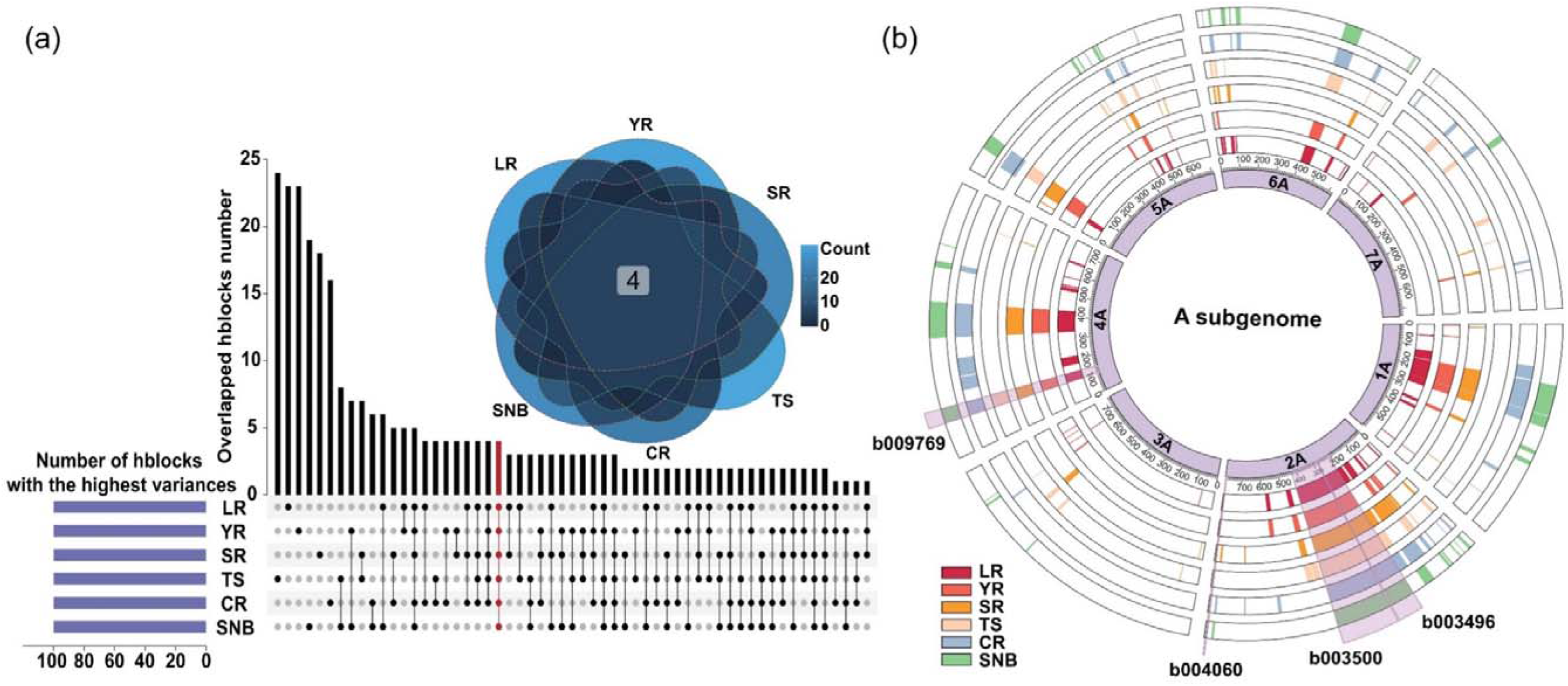
Pleiotropic haploblocks (hblocks) identified from the top 100 blocks with the highest haplotype variances for each respective disease. **(a)** Upset and Venn diagram showing the number of overlaps responsible for different diseases. LR, leaf rust; YR, yellow rust; SR, stem rust; TS, tan spot; CR, crown rot; SNB, Septoria nodorum blotch. The red indicates the haploblocks identified for all the six diseases. **(b)** Circos plot showing the chromosome distributions of the top 100 haploblocks for each disease. Only A subgenome is displayed here. The four haploblocks related to six diseases are highlighted with purple shadows.

The four haploblocks, b003496, b003500, b004060 and b009769 with high resistance variances for six diseases were located on chromosome segments 2AS (209.71-276.25 Mb), 2A (279.8-427.97 Mb), 2AL (780.97-784.79 Mb), and 4AS (100.09-139.91 Mb), respectively (Table S12). Previously reported QTLs from studies on individual diseases were positioned in the context of the haploblocks reported in this study. Meanwhile, candidate genes likely involved in diverse defense mechanisms were predicted, which could contribute to the resistance of each haploblock for various pathogens (Table S13). We identified all haplotype variants present in the Vavilov diversity wheat panel, and phenotypic effects contributed by different haplotypes of the four haploblocks to six diseases were displayed (Figure S20). Marked differences were observed for most disease responses between wheat accessions carrying haplotype variants of the highest and lowest effects (Figure S20). Further, four superior haplotypes at b004060 and three superior haplotypes at b009769 showed favourable effects for all six diseases (Figure S21), suggesting these haplotypes could be utilised to improve MDR in breeding programs. Interestingly, the three wheat accessions carrying the haplotypes were all landraces from Asia (Table S14).

In conclusion, we have provided a haploblock-based navigation catalogue with novel forms of resistance, resulting in the discoveries of genetic regions consistently explaining genetic variation across environments for rust resistance and key haploblocks underlying MDR in the Vavilov wheat panel.

### Assessing genetic potential of multiple disease resistance by superior haplotype stacking

To explore the potential of breeding for MDR, we first investigated and screened individual accessions of Vavilov wheat diversity panel according to Ri values which is appropriate to screen resistance (Riaz et al., 2018) and the accessions with below 0 values had a high level of resistance across different environments, whereas values above 0 implied susceptibility (Figure S8). Based on Ri of the six diseases, the 295 diverse wheat accessions clustered into three groups (Figure 22a), where cluster 3 had higher resistance to LR and YR but was more susceptible to SNB compared to cluster 1 (Figure 22b & 22c). In total, there were 19 out of 295 wheat entries screened with high resistance to at least five different diseases, and most of them were landraces representing the highest degree of MDR in the wheat panel (Table S15).

Taking leaf rust as an example, contrasting genomic components present by genome-wide haplotype effects were observed between the most LR resistant and susceptible wheat accessions, that is, the resistant wheat accessions carried more superior haplotypes, whereas the susceptible accessions carried more haplotypes with negative effects (Figure 3a). Further using all Vavilov wheat accessions and the top 100 LR haploblocks, we showed that cumulative detrimental haplotype counts decreased in frequency incrementally with LR resistance (Figure 3b). These results supported the methodology based on local GEBVs and suggested that superior haplotype stacking could be an effective strategy to improve disease resistance.

**Figure 3.**
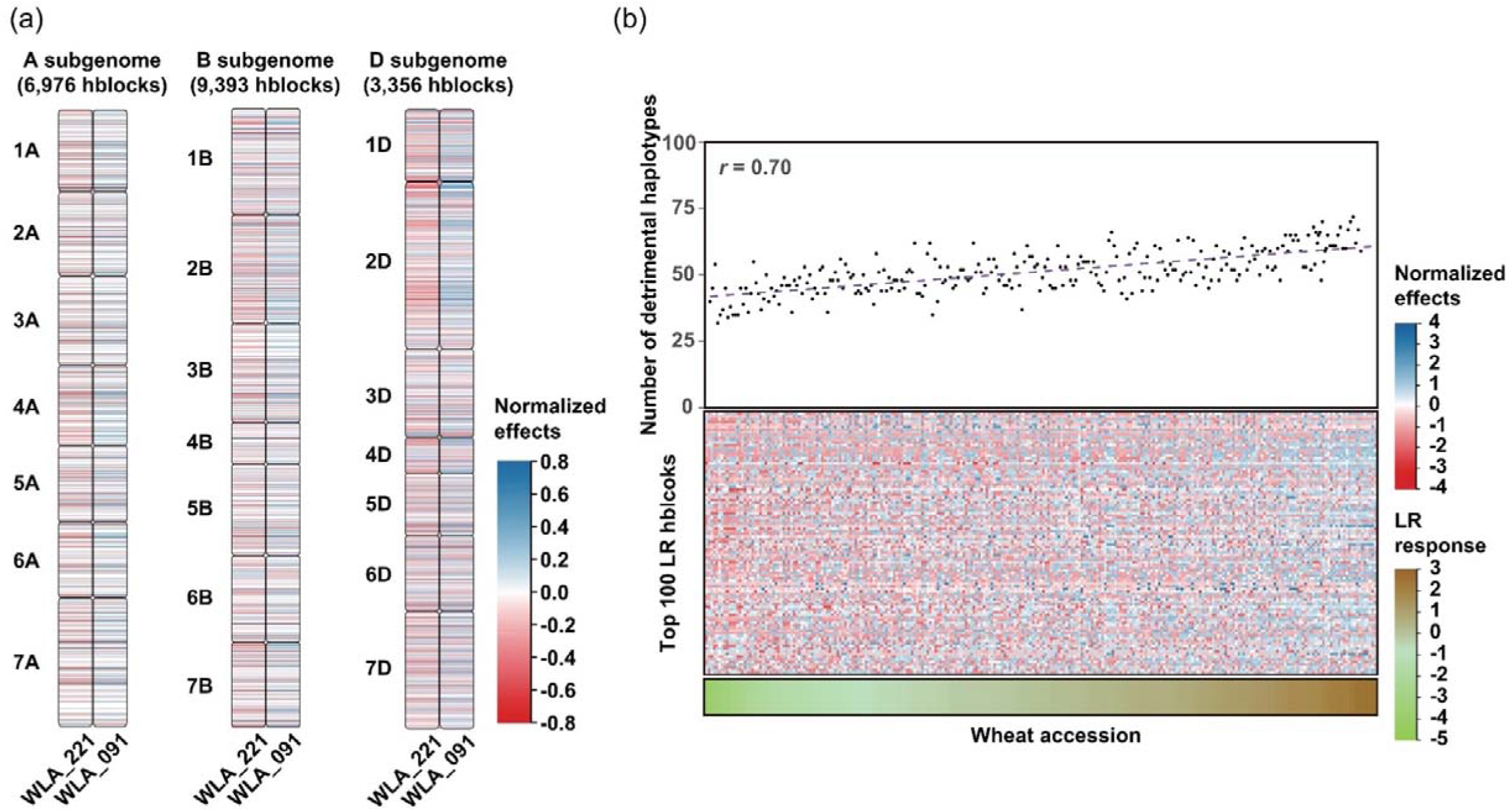
Haplotype stacking is an effective avenue for resistance breeding. **(a)** Comparison between the most resistant (WLA_221) and susceptible (WLA_091) wheat accession for leaf rust (LR) in the Vavilov panel, where red represents favourable haplotypes and blue represents detrimental haplotypes. **(b)** Distribution of the top 100 LR haploblocks along with trait performance in all Vavilov wheat accessions. *r*-value is the Pearson correlation of the detrimental haplotype counts to LR response for each wheat accession. Red represents favourable haplotypes, whereas blue represents detrimental haplotypes. Increasing green represents increasing LR resistance.

Based on a simulation approach creating *in silico* wheat genotypes that can stack different numbers of superior haplotypes at haploblocks weighted based on their estimated variances, we predicted changes in resistance levels for all six diseases by stacking the most favourable haplotypes for each individual disease (Figure 4a). Ten wheat accessions with the highest resistance of each respective disease were used as a basis for simulation, and *in silico* genotypes were generated from these selected accessions by replacing their original haplotypes with the most favourable haplotypes. As expected, haplotype stacking could improve resistance potential substantially in the ten most resistant wheat accessions for the given disease, but the indirect effects on the other five diseases varied greatly. For example, stacking the haplotypes most favourable to LR led to the improvement of LR resistance potential but simultaneously resulted in decreased resistance to SNB. In this example, replacing all 19,725 genome-wide haploblocks increased LR resistance by up to three-fold, but SNB inadvertently decreased by nearly two-fold (Figure 4a). These results were consistent with the negative correlation between LR and SNB (Figure 1a). Interestingly, stacking the most favourable haplotypes for YR indirectly improved resistance to both LR and SR. In this case, stacking at all the desirable haploblocks increased YR, SR and LR resistance potential by up to three-, two-, and one-fold, respectively (Figure 4a).

**Figure 4.**
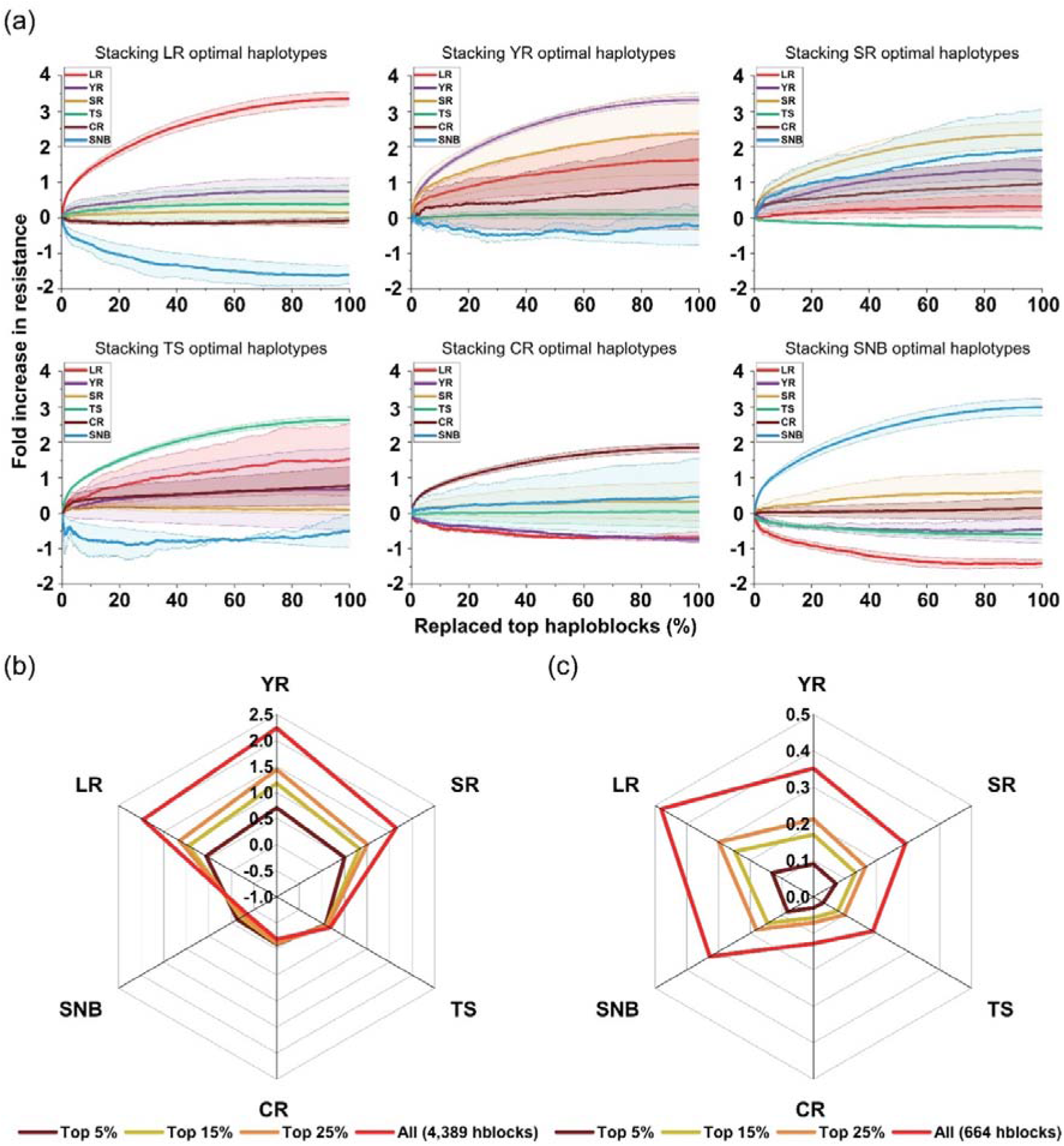
Genetic potential of multiple disease resistance (MDR) revealed by stacking superior haplotypes. **(a)** Resistance predictions evaluating effects of stacking optimal haplotypes for each individual disease on the six diseases. LR, leaf rust; YR, yellow rust; SR, stem rust; TS, tan spot; CR, crown rot; SNB, Septoria nodorum blotch. The ten wheat accessions with the highest resistance of each respective disease were used as a basis for simulation. *In silico* genotypes were generated from these selected cultivars by exchanging the original haplotypes by the most favourable haplotypes at the haploblocks. Relative changes to genomic estimated breeding values (GEBV) were calculated. The solid lines indicate the mean of predicted changes, and standard errors are displayed by colored shadows with dashed lines. **(b)** Radar plot illustrating the predicted mean fold increase of MDR by stacking three-rust superior haplotypes. **(c)** Radar plot illustrating the predicted mean fold increase of MDR by stacking six-disease superior haplotypes. Nineteen wheat accessions screened with high resistance to at least five diseases were used as a basis for simulation.

To investigate direct improvement of MDR through haplotype stacking, we selected the haplotypes with positive effects on multiple diseases. In total, 4,389 and 664 haploblocks were associated with beneficial effects for the three rusts and six diseases, respectively (Figure S23; Table S16-S17). As expected, *in silico* lines created through continuously stacking three-rust optimal haplotypes incrementally increased resistance to the rusts compared with the 19 real wheat accessions with the highest MDR, although resistance for other three diseases (TS, CR and SNB) hardly changed. On the other hand, stacking haplotypes most favourable to six diseases increased MDR for all diseases (Figure S24 & 4b). Replacing only 5–25% of the haploblocks (i.e. 33–166) in the 19 best MDR wheat accessions could simultaneously increase six-disease resistance by nearly 5–30% on average, while accumulation of the optimal haplotypes at all 664 haploblocks was predicted to increase resistance to LR, YR, SR, TS, CR and SNB by up to 48.1, 35.2, 29.1, 18.8, 12.8 and 32.8%, respectively, compared to the most resistant accessions in the Vavilov wheat diversity panel (Figure 4b).

To conclude, through simulation we demonstrate stacking optimal haplotypes for one disease could improve resistance for positively correlated diseases, but not negatively correlated diseases, and there was considerable potential to develop MDR wheat cultivars by stacking multi-disease resistant haplotypes.

## Discussion

To the best of our knowledge, this is the first study to estimate global and local genetic correlations for resistance to major fungal diseases in wheat. We established a haplotype catalogue based on genome-wide haploblocks with novel forms of resistance, resulting in the discovery of multi-disease resistance haplotypes in the Vavilov collection. The breeding potential of available single-and multi-disease resistance was predicted through haplotype stacking simulations. The diversity within the Vavilov wheat panel is anticipated to play an important role in contributing to haplotype-based breeding approaches for MDR.

### Vavilov wheat datasets help unravel the neglected genetic relationships of resistance associated with six major wheat diseases

Although genetic correlations between complex traits have been extensively investigated in human and animal genetics, insight into genetic relationships of disease resistance traits is quite limited in plants (van Rheenen et al., 2019). The polygenic architecture of resistance traits for the six wheat fungal diseases is well established and variation in these traits is underpinned by many additive major and minor genes. However, pathogen lifestyles and resistance mechanisms of plants vary among rust diseases, TS/SNB, and CR (Figueroa et al., 2018). In the Vavilov panel, positive correlations were observed between the resistance of the three rusts, probably due to similar immune mechanisms and underlying resistance genes of plants (Norman et al., 2023). The three rusts do not always occur in the field concurrently, since their infection depends on the climatic conditions that are different during plant growth. However, positive genetic correlations imply the possibility of host resistance against various rust pathogens throughout wheat growth stages.

Interestingly, CR was positively correlated with SR, and both causal pathogens have the ability to infect the stem of wheat plants, where similar defence mechanisms might occur (Figueroa et al., 2018). The negative correlation between necrotrophic SNB and biotrophic LR/YR might be explained by the likelihood of opposing resistance mechanisms (Friesen and Faris, 2021; Wiesner-Hanks and Nelson, 2016). In the case of SNB, a dominant host susceptibility (*S*) gene can lead to programmed cell death (PCD), thereby facilitating pathogen infection. However, this process also acts in a similar manner as a hypersensitive response against biotrophic pathogens, which restricts pathogen infection (Wiesner-Hanks and Nelson, 2016). Concurrently, other genetic factors add a layer of complexity to interactions between necrotrophic and biotrophic diseases. For example, pattern-triggered immunity (PTI) aids plants to defend the infections of multiple pathogens (Lu and Tsuda, 2021), which could play a dominant role for a positive correlation between TS and YR. One QTL resistant to TS coincided with *Lr46*/*Yr29*/*Sr58*, suggesting a haplotype associated with *Lr46*/*Yr29*/*Sr58* could also contribute to TS resistance (Singh et al., 2019). Interestingly, a significant negative correlation was observed between TS and SNB. Possible explanations could be (1) previous studies reported some race-nonspecific resistance QTL acting upstream of the ToxA-*Tsn1* interaction, which might cause the different resistance of host to TS and SNB (Hu et al., 2019); or (2) SNB and TS have distinct genetic controls acting at various growth stages, whereby in previous studies, SNB was investigated using specific pathogen strain in glasshouse at seedling stage, while TS was in field, where variation in pathogen population was less controlled or unique genetic mechanism associated with adult-plant resistance (Dinglasan et al., 2019; Phan et al., 2018).

Our study analysed local genetic correlations by haploblock-based calculations, which could hint at common genomic regions regulating resistance to wheat diseases. Similar genome-wide patterns were observed for genetic correlation estimates from the CS method (Figure S15), implying the true existence of genomic regions that are genetically correlated. A study in cattle also showed that the marker effects were not biased by LD, that is, for the LD pruned markers, no substantial differences were found for the genome-wide pattern of the *r* estimates (Olasege et al., 2022). Some genes might be involved in multiple traits that are not correlated, leading to identification of significantly correlated resistance haploblocks for uncorrelated diseases (Figure 1). However, in general, higher global genetic correlations could be attributed to higher local genetic correlations, of which more shared ‘driver’ haploblocks were detected between trait pairs. Studies on human diseases agree with these findings, which suggested that overall genetic correlations were underpinned by average correlations at local genomic regions (Guo et al., 2021; Werme et al., 2022).

### Genetic basis of multiple disease resistance revealed by multi-trait haplotype characterisation

GWAS is a commonly-used genetics approach to unravel the rich genetic basis of complex wheat disease resistance traits (Hamblin et al., 2011). In this study, we adopted a haplotype-based mapping approach to create a resistance catalogue showcasing the genetic effect of each haplotype in the wheat genome for resistance to six major diseases across diverse environments. Distinct from the single marker approach used in traditional GWAS studies, we estimated the genetic effects of chromosome segments in pronounced LD. The resultant chromosome segments relevant to multiple traits could be more desirable to be used in breeding, since pyramiding alleles for specific traits can be more costly and time-consuming, while stacking of a large number of loci is nearly impossible due to excessive generations required (Hafeez et al., 2021). Chromosome segments defined by LD can be more easily manipulated in a breeding program because the group of alleles on a haplotype is not prone to losing linkage with the trait during crossing and recombination. Meanwhile, a haploblock might also harbour molecule modules integrating multiple causal genes or alleles with holistic effects (Song et al., 2023).

Tapping into the haploblock-based resistance catalogue, 26 genetic regions associated with consistent resistance variation for LR (13), YR (1), and SR (12) were considered novel (Table S7-S9). Another 26 were co-located with the reported QTL in previous studies and this builds confidence towards their true genetic effects. Different from relatively simple candidate gene prediction in GWAS, genetic intervals of the haploblocks detected here tend to be larger, which can complicate the discovery of causal genes and future isolation. However, this can be further resolved by increasing *r* parameters in defining the LD blocks when exploring local LD in ‘fine-mapping’. Nevertheless, haplotype markers with considerable effects within the haploblock regions can be used to assist the straightforward selection of favourable haplotypes, facilitating the utilisation of these identified haploblocks in a range of breeding scenarios or strategies.

The discovery of pleiotropic multi-disease resistance haploblocks with high variances suggests there may be shared genetic controls for MDR and, thus, warrants further investigation. Of the four haploblocks associated with all six diseases, three co-located with previously reported QTL for some of the individual diseases, but not to all six diseases. Interestingly, b009769 (100.09-139.91 Mb, 4AS) has not previously been associated with any of the diseases (Table S12), implying that haplotype-based mapping could effectively identify chromosomal segments for MDR. Within the four haploblocks for MDR, a number of candidate gene were present likely involved in various resistance pathways against biotrophic and/or necrotrophic diseases, including signal transduction, PCD, sugar metabolism, cell wall thickness, phytohormone regulation, and detoxification (Table S13) (Bogacki et al., 2008; Friesen and Faris, 2021; Lu and Tsuda, 2021). Resistance gene clusters were also discovered, such as *TraesCS2A03G1354300*, *TraesCS2A03G1354400*, and *TraesCS2A03G1354500* encoded for a dirigent protein and two NBS-LRR proteins, respectively. TaDIR-B1, a dirigent protein, was recently characterised to participate in CR resistance via influencing lignin content of plant stem, supporting the potential role of *TraesCS2A03G1354300* (Yang et al., 2021). MDR might be attributed to the combinations of these candidate genes that offer diverse defence signalling or pathways to protect against multiple pathogens (Hafeez et al., 2021). In a recent study, a haplotype-based selection strategy led to the coordinated improvement of plant architecture, yield, and nitrogen-use efficiency by selecting for a haploblock harbouring a molecule module, *Rht-B1b–EamA-B–ZnF-B* involved in various hormone pathways in wheat (Song et al., 2023). As most cloned *R* genes are pathogen-specific receptor proteins (Sánchez-Martín and Keller, 2021), selection or introgression of favourable haplotypes capturing multiple alleles or holistic genetic effects, rather than a single allele, could assist the improvement of MDR.

### Exploiting genetic potential for multiple disease resistance breeding

While the most LR resistant wheat accession in the panel carries many superior haplotypes for LR (i.e. WLA_221), the most susceptible accession in the panel (WLA_091) also carries favourable haplotypes, some of which are absent in WLA_221 (Figure 3a). Therefore, both accessions provide complementary donor sources to enrich breeding progenies with beneficial haplotypes. Notably, WLA_221 carries detrimental haplotypes at over 25 out of the top 100 LR haploblocks with the highest variances (Figure 3b), which highlights the potential for further genetic improvement by replacing these deleterious haplotypes through breeding. Technologies that enable induced targeted recombination have the potential to facilitate favourable haplotype stacking (Ru and Bernardo, 2019), but the indirect effects on other diseases need to be considered to develop new cultivars incorporating MDR. The optimal strategy to improve multiple rust resistance appears to be direct selection for YR superior haplotypes (Figure 4a). SR haplotype stacking was inadvertently beneficial for SNB, whereas stacking LR superior haplotype significantly decreased SNB resistance, vice versa, indicating a potential trade-off for selection of LR and SNB resistance in breeding programs. These results are largely consistent with the correlations estimated between disease traits, and the presence of ‘driver’ or ‘antagonizing’ haploblocks also supports the reasons why stacking optimal haplotypes for one disease can impact resistance levels for other diseases. A lack of knowledge of MDR could impede the development of resistant cultivars and unintentionally cause epidemics of the diseases that are genetically correlated but have been overlooked.

In the analysis of the 19 wheat accessions with the highest level of MDR in the Vavilov panel, we demonstrate there is still large genetic potential to generate ‘super’ wheat cultivars with high MDR by stacking different haplotypes associated with multi-disease resistance (Figure 4b). Haplotype-based genomic selection algorithms, such as genetic algorithm (Kemper et al., 2012) and maximum haploid breeding value calculation (Müller et al., 2018), can be used to identify parental lines that maximise diversity for resistance haplotypes. Following selection of parents, simulation can help identify the most optimal crossing path to combine the target haplotypes into elite cultivars (Villiers et al., 2023), with the potential to accelerate the crossing and population development through speed breeding (Watson et al., 2018). A current limitation of this approach is the limited understanding of G × E × P (genotype × environment × pathogen) interactions that influence disease response (Pariaud et al., 2013). Large-scale, high quality disease response datasets for multiple pathotypes and diverse field environments are required to estimate these effects at the haploblock level. Overall, the resistance catalogue established in this study provides a first step and valuable resource for the wheat research and breeding community to implement haplotype stacking approaches and accelerate the genetic gain for MDR in wheat.

## Experimental procedures

### Plant materials

The Vavilov wheat diversity panel was used in this study. The panel contained 295 diverse wheat accessions originally sourced from the N. I. Vavilov Institute of Plant Genetic Resources (VIR) (Riaz et al., 2017). The panel comprised 136 landraces, 32 cultivars, 10 breeding lines and 118 accessions with unknown cultivation status, which were collected between 1922 to 1990 (Table S1). A total of 206 entries had known origin information, originating from 28 countries, spanning North America (4), South America (2), Africa (6), Europe (69), and Asia (125).

### Genotyping and integrated marker map construction

The Vavilov wheat diversity panel was genotyped by 90K SNP array, single-row DArT SNP and silico-DArTseq platforms, which returned a total of 51,852, 62,273, and 57,287 markers, respectively. Flanking sequences of each marker were blasted on IWGSC RefSeq v2.1 (Zhu et al., 2021) using TBtools software (Chen et al., 2020) to determine marker physical positions by the best hit results (*E*-value < 1×10^-5^). After discarding the duplicates and markers with >20% missing data and a minor allele frequency < 5%, 34,899 high-quality markers were finally retained and ordered onto an integrated marker map according to their physical positions.

### Population and diversity analyses

To identify population structure and diversity of the Vavilov wheat panel, different approaches were carried out, including Bayesian clustering in Structure v2.3.4 software (https://web.stanford.edu/group/pritchardlab/structure_software/), Ward’s hierarchical clustering and Kinship analysis in R package ‘GAPIT v3.0’ (Wang and Zhang, 2021), as well as PCA and Neighbor-Joining tree analysis in Tassel v5.0 software (Bradbury et al., 2007). To explore average LD decay of the panel, pairwise LD was estimated using the squared allele frequency correlation (*r*^2^) among markers by Tassel v5.0, followed by locally weighted regression (LOESS) of pairwise LD against physical distances between markers and curve smoothing performed in R v4.0.2 with function ‘loess.smooth’.

### Phenotypic analyses of six different fungal diseases

Phenotypic data for disease responses to LR, YR, SR, TS, SNB and CR were collected from 22 experiments (Table S2). LR and YR field trials were conducted and described in our previous studies (Jambuthenne et al., 2022; Riaz et al., 2018) to phenotype the Vavilov wheat accessions in Queensland (QLD) over three consecutive cropping seasons (2014–2016), which are hereafter named as LR_14QLD, LR_15QLD, LR_16QLD, YR_14QLD, YR_15QLD, YR_16QLD, respectively. In addition, eight field experiments were conducted to evaluate YR resistance in Ethiopia, i.e. Meraro (2018, 2019, 2021), Kulumsa (2018, 2019, 2021), and Bekoji (2019, 2021), by pathogen natural infection with the predominant races Pst11 and Pst16 in the seasons (https://agro.au.dk/forskning/internationale-platforme/wheatrust/). The experiments are referred here as YR_18MR, YR_19MR, YR_21MR, YR_18KU, YR_19KU, YR_21KU, YR_19BK, YR_21BK, respectively. Resistance to SR was assessed through four field trials in Debrezeit in 2018, 2019 (two growth seasons), and 2021 using a mixture of *Pgt* isolates (TKKTF, TKTTF, TRTTF, TTKSK, TTKTT, and TTRTF). The four experiments are hereafter named SR_18DZ, SR_19DZ, SR_19DZo, and SR_21DZ, respectively. Phenotyping for TS was previously described in our study (Dinglasan et al., 2019), wherein two field experiments were carried out and referred here as TS_15QLD and TS_16QLD, respectively. One field trial (named as CR_18QLD) was conducted in 2018 for CR resistance in a CR screening nursery established in QLD (Alahmad et al., 2020), and a pure culture of *F. pseudograminearum* isolates was collected (Alahmad et al., 2018) and used to infect wheat accessions. Seedling response for SNB (isolate SN15) (Phan et al., 2018) was used to represent the SNB resistance level of each Vavilov wheat accession and hereafter named as SNB_18WA. Collectively, these phenotypic data were from the pathotypes that were highly virulent and widespread across the Australian or Ethiopian wheat-growing regions. To enable all the phenotypic datasets with the same scale, coefficients of infection (CI, 0–100) for the three rust diseases assessed at the adult plant stage were converted to the scale 0–9 (Jambuthenne et al., 2022; Riaz et al., 2018).

To effectively simplify or reduce the dimensions of the various phenotypic datasets, three parameters, i.e. relative disease index (Ri), best linear unbiased estimate (BLUE) and eigenvector principal component 1 (PC1) explaining the most of phenotypic variances across environments, were calculated in each wheat accession in each disease. Ri was calculated for each wheat accession in each disease as the following equation (Jambuthenne et al., 2022; Riaz et al., 2018):

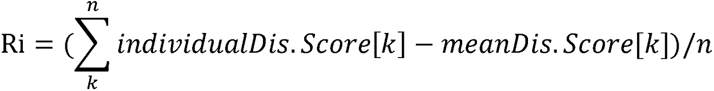

where *n* is the total experiment number, *k* is the individual experiment number. Ri below 0 indicates a relatively high level of resistance, while value above 0 indicates susceptibility.

BLUE value of each wheat accession was estimated by fitting the following mixed linear model in R package ‘lme4’:

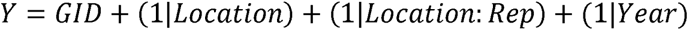

where *Y* is disease response data, the parentheses indicate random effects, ‘1|’ indicates groups, ‘:’ represents interactions. *GID* represents all wheat genotypes used, *Location* and *Year* indicates experimental location and year, respectively, and *Rep* is the replications in one experiment.

R package ‘FactoMineR’ was used to impute missing phenotypic values through expectation maximization method in PCA, followed by estimation of percentage of explained variance for each PC and calculation of PC1 (http://factominer.free.fr/index.html).

### Trait Pearson correlation and global genetic correlation

Ri, BLUE and PC1 were each used for calculations of Pearson correlation and global genetic correlation between six different disease traits. For 22 individual environments, the correlation estimations and PCA of all the experiments were carried out. Pearson correlation analysis and PCA was performed in R packages ‘corrplot’ and ‘FactoMineR’, respectively. Genome-based restricted maximum likelihood (GREML) bivariate model was performed in GCTA software (Yang et al., 2011) to calculate pair-wise global genetic correlations between traits.

### Genome-wide haploblock construction

Genome-wide LD block (haploblock) construction was performed using an algorithm implemented in the R package ‘SelectionTools’ (http://population-genetics.uni-giessen.de/~software/). First, pairwise *r^2^* values between markers were calculated across the 21 wheat chromosomes, followed by selection of adjacent pairs on each chromosome with the highest LD among all pairs. If the *r*^2^ of a pair exceeded the threshold, these markers were defined as a new haploblock. In the next step, flanking markers on each side of the haploblock were considered and added to the same block if their pairwise *r^2^* values with the respective outer markers in the LD block also exceeded the threshold. At the same time, the tolerance parameter (*t*) allowed *t* markers present within a block, that is, if a flanking marker did not fulfill the LD threshold, the block was still extended if at least *t* adjacent flanking markers fulfill the LD grouping threshold. The haploblock was completed if more than *t* flanking markers had a lower LD than the threshold. This procedure was repeated until all markers were assigned to haploblocks. Markers not in LD with any others were assigned to individual haploblocks. Different *r^2^* (0.5, 0.6, 0.7, 0.8, 0.9) and *t* value (1, 2, 3) combinations were tested for above procedures. Finally, genome-wide haploblocks were constructed by adopting *r*^2^ = 0.5 and *t* = 3. Physical position of each haploblock on chromosome and block size were defined by the included markers (Table S4).

### Local genetic correlation by haploblock-based calculation

First, we estimated single marker effects for each disease trait, using a ridge-regression best linear unbiased prediction (rrBLUP) model fitting all markers simultaneously (Endelman, 2011). We then correlated marker genetic effects within each haploblock (see above) harbouring >3 adjacent markers in LD between all 15 pairs of disease traits. Pearson correlation coefficient (*r*) was calculated, which was formulated as below:

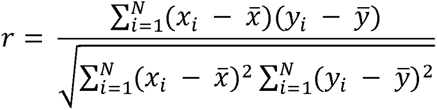

where *N* is the number of markers in each haploblock, *x_i_* and *y_i_* is the marker effect of each trait in a trait pair, and *x̅* and *y̅* is the mean value of *x* and *y*, respectively. Only *r* estimates with Bonferroni-corrected *P*-value < 0.05 were considered as significant. Linear regression was performed in R package ‘ggplot2’ between mean correlations of significant haploblocks and global genetic correlations (see above) using all 15 trait pairs. Finally, the ‘driver’ and ‘antagonizing’ haploblocks were defined as the haploblocks driving or antagonizing the global genetic correlations, respectively (Olasege et al., 2022). If the global genetic correlation between two traits is positive/negative, the ‘driver’ regions are the haploblocks with significant positive/negative correlations, while the ‘antagonizing’ haploblocks are with significant negative/positive correlations. We also used correlation scan, a recently developed method (Olasege et al., 2022) to identify local genomic correlations in livestock, to compare with the haploblock-based method.

### Local GEBV calculation and genome-wide resistance catalogue development

Local genomic estimated breeding values (local GEBVs) for haplotypes within each chromosomal block were calculated and a haplotype-based mapping approach was employed to identify chromosome segments (i.e. haploblocks) with a high impact on the respective disease trait in the panel. The local GEBV method simultaneously discovers favourable haplotypes with strong effects on traits while accounting for selection signatures revealed by pronounced genome-wide LD (Voss-Fels et al., 2019). Briefly, rrBLUP marker effects and the constructed genome-wide haploblocks (see above) were used in this analysis. Since the rrBLUP model fits all markers simultaneously, the procedure does not rely on single-marker *P-value* statistics, which is prone to overestimation of effects. Next, the haplotypes of each haploblock were identified in the Vavilov diversity population, and the haplotype effects, also referred as ‘local GEBVs’ were calculated by the sum of the single marker effects. Finally, the variances of the local GEBVs were calculated in each haploblock, and a higher variance indicates the haploblock is more likely to be associated with the target trait (Voss-Fels et al., 2019).

### Analyses of multi-disease resistance haploblocks with consistently high variances across environments

Based on the haploblock-based resistance catalogue above, we considered the top 100 (i.e. top 0.5%) haploblocks with the highest variances as trait-relevant genetic regions (Voss-Fels et al., 2019). Consistent haploblocks identified in all field experiments were obtained for each of the three rust diseases. These haploblocks were compared with previously mapped QTL related to rust resistance. In terms of multiple diseases, PC1 values for each of the six disease traits were used to estimate whole-genome haploblock variances. Pleiotropic haploblocks were considered those that occurred in the top 100 haploblocks for at least two diseases. Candidate genes situated in the pleiotropic haploblocks were examined based on gene annotations obtained from IWGSC RefSeq v2.1 (Zhu et al., 2021). The genes annotated as being involved in potential disease resistance pathways were listed as the candidate genes underlying the identified haploblocks.

### Exploring the potential of haplotype stacking for single and multi-disease resistance

The whole genomic components of the most LR resistant and susceptible wheat accessions were compared using haplotype effects (local GEBVs). Further, using all Vavilov wheat accessions and the top 100 LR haploblocks, cumulative effects of haplotypes was evaluated by linear regression analysis between the detrimental haplotype counts to LR responses of each wheat accession.

The effects of stacking optimal haplotypes for each disease was evaluated to predict the change in resistance levels for all six diseases. Ten wheat accessions with the highest resistance of each respective disease were used as a basis for simulation. *In silico* genotypes were generated from these selected accessions by exchanging their original haplotypes by the most favourable haplotypes at the haploblocks. The haploblocks were weighted by their variances on the given disease to be replaced from the major to minor. Each time, 0.1% haploblocks were replaced until all 19,725 haploblocks were replaced in the selected real accessions. The fold change in resistance for each disease was denoted by GEBV comparisons of each pair of *in silico* and real genotypes, as the following calculation:

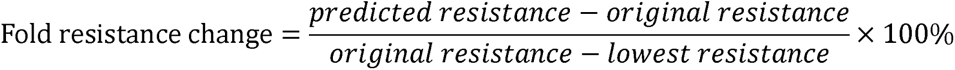

where *predicted and original resistance* is denoted by the sum of GEBVs after and before replacement of haplotypes, respectively, and the value of *lowest resistance* differs in each disease. The analysis was performed for each of the six diseases.

To investigate multi-disease resistance potential, we simulated stacking of only the optimal haplotypes favourable for multiple diseases. The haploblocks and haplotypes that have beneficial effects on three rusts and six diseases were each identified and targeted for stacking. Nineteen wheat accessions screened with high resistance to at least five diseases were used as a basis for simulation. The most favourable multi-disease haplotype was determined at each pre-selected haploblock by maximising the sum of effects on each disease. Following the same simulation approach above, the potential of multi-disease resistance improvement was assessed by exchange of the original haplotype by the most favourable haplotype at each of pre-selected multi-disease haploblocks. The analysis was performed for each three-rust and six-disease breeding scenario.

## Acknowledgements

We are grateful to Dilani Jambuthenne, Adnan Riaz and Kar-Chun Tan for sharing disease response data for the Vavilov diversity panel. This research was supported by the Grains Research and Development Corporation (CSP2304-013RTX). JT received a scholarship from the China Scholarship Council (202203250009). LH was supported through an ARC Future Fellowship (FT220100350).

## Author contributions

JT analysed the data and wrote the manuscript. ZT and SA contributed phenotypic data. LH, SP, ED and BH supervised the research project and critically reviewed and revised the manuscript. All of the authors read and approved the final version of the manuscript.

## Supporting information

**Figure S1** Integrated marker map with 90K, DArT SNP, and DArTsilico (DArTseq) markers.

**Figure S2** Marker allele distributions of integrated map on each chromosome in 295 Vavilov wheat accessions.

**Figure S3** Population structure of the 295 Vavilov wheat accessions revealed by Ward’s hierarchical clustering, Kinship, and STRUCTURE software analyses.

**Figure S4** Three subpopulations are classified in the Vavilov wheat panel.

**Figure S5** Linkage disequilibrium (LD) decay over physical distances on the A, B, D subgenome, and the whole genome.

**Figure S6** Phenotypic variations and distributions of 22 individual experiments on disease responses to leaf rust (LR), yellow rust (YR), stem rust (SR), tan spot (TS), crown rot (CR) and Septoria nodorum blotch (SNB).

**Figure S7** Principal component analysis (PCA) of phenotypic datasets of leaf rust (LR), yellow rust (YR), stem rust (SR), and tan spot (TS).

**Figure S8** Distributions of the three phenotypic parameters representing the resistance level of each wheat accession.

**Figure S9** Relationships between six wheat major diseases evaluated by three different phenotypic parameters, relative disease index (Ri), eigenvector principal component 1 (PC1), and best linear unbiased estimate (BLUE).

**Figure S10** Relationships between 22 individual experiments on disease responses to leaf rust (LR), yellow rust (YR), stem rust (SR), tan spot (TS), crown rot (CR) and Septoria nodorum blotch (SNB).

**Figure S11** Genetic correlations among 22 individual experiments on disease responses to leaf rust (LR), yellow rust (YR), stem rust (SR), tan spot (TS), crown rot (CR) and Septoria nodorum blotch (SNB).

**Figure S12** Genome-wide distribution and size of defined haploblocks.

**Figure S13** Distribution patterns of haploblocks across all the 21 wheat chromosomes.

**Figure S14** Genome-wide plot of the local genetic correlation estimates for 15 disease trait pairs.

**Figure S15** Comparison of local genetic correlation calculations between haploblock-based method and correlation scan (CS).

**Figure S16** Significant driving and antagonizing haploblocks along with genetic correlations between all 15 trait pairs.

**Figure S17** Haploblocks contribute to high yellow rust (YR) correlations within and between Queensland (QLD) and Ethiopia environments.

**Figure S18** Whole-genome haploblock variances for the six diseases using eigenvector principal component 1 (PC1).

**Figure S19** Circos plot showing the chromosome distributions of the top 100 haploblocks on B and D subgenomes for each of the six diseases.

**Figure S20** Phenotypic effects contributed by haplotypes of the four haploblocks, b003496, b003500, b004060 and b009769 related to six diseases.

**Figure S21** Superior haplotypes of b004060 and b009769 with positive effects to improve resistance levels of the six diseases.

**Figure S22** Screening of Vavilov wheat accessions based on relative disease index (Ri) of six diseases.

**Figure S23** Number of haploblocks and corresponding haplotypes that can have beneficial effects on three rusts and six diseases.

**Figure S24** Resistance predictions evaluating effects of stacking the most beneficial haplotypes to three rusts and six diseases on multiple disease resistance (MDR).

**Table S1** Passport information of 295 Vavilov wheat accessions used in this study.

**Table S2** Descriptive statistics on the integrated marker map.

**Table S3** Detailed information on the phenotypic disease datasets investigated in this study.

**Table S4** Genome-wide 19,725 haploblocks constructed in the Vavilov panel.

**Table S5** Local genetic correlation estimates from haploblock-based calculations for 15 disease trait pairs.

**Table S6** Resistance catalogue based on genome-wide haploblocks showcasing corresponding variances for six disease traits among 295 Vavilov wheat accessions measured across 22 individual environments.

**Table S7** Consistent haploblocks identified in all environments for leaf rust (LR).

**Table S8** Consistent haploblocks identified in all environments for stripe rust (YR).

**Table S9** Consistent haploblocks identified in all environments for stem rust (SR).

**Table S10** Consistent haploblocks of stripe rust (YR) identified in both Queensland (QLD) and Ethiopia environments.

**Table S11** Pleiotropic haploblocks with high impact on at least two different diseases.

**Table S12** Four haploblocks with high impact on six wheat diseases, including leaf rust (LR), yellow rust (YR), stem rust (SR), tan spot (TS), crown rot (CR) and Septoria nodorum blotch (SNB).

**Table S13** Predicted causal genes underlying the four pleiotropic haploblocks relevant to six diseases.

**Table S14** The Vavilov wheat accessions carrying with the superior haplotypes beneficial to six diseases for b004060 and b009769.

**Table S15** The Vavilov wheat accessions with high resistance to at least five diseases screened by relative disease index (Ri).

**Table S16** The haploblocks and corresponding haplotypes most favourable to three rust diseases.

**Table S17** The haploblocks and corresponding haplotypes most favourable to six diseases.

