## Supplementary material for "Stacking haplotypes from the Vavilov wheat collection to accelerate breeding for multiple disease resistance": Supplementary figures.pdf

**Running title:** Genetic potential of multi-disease resistance in wheat

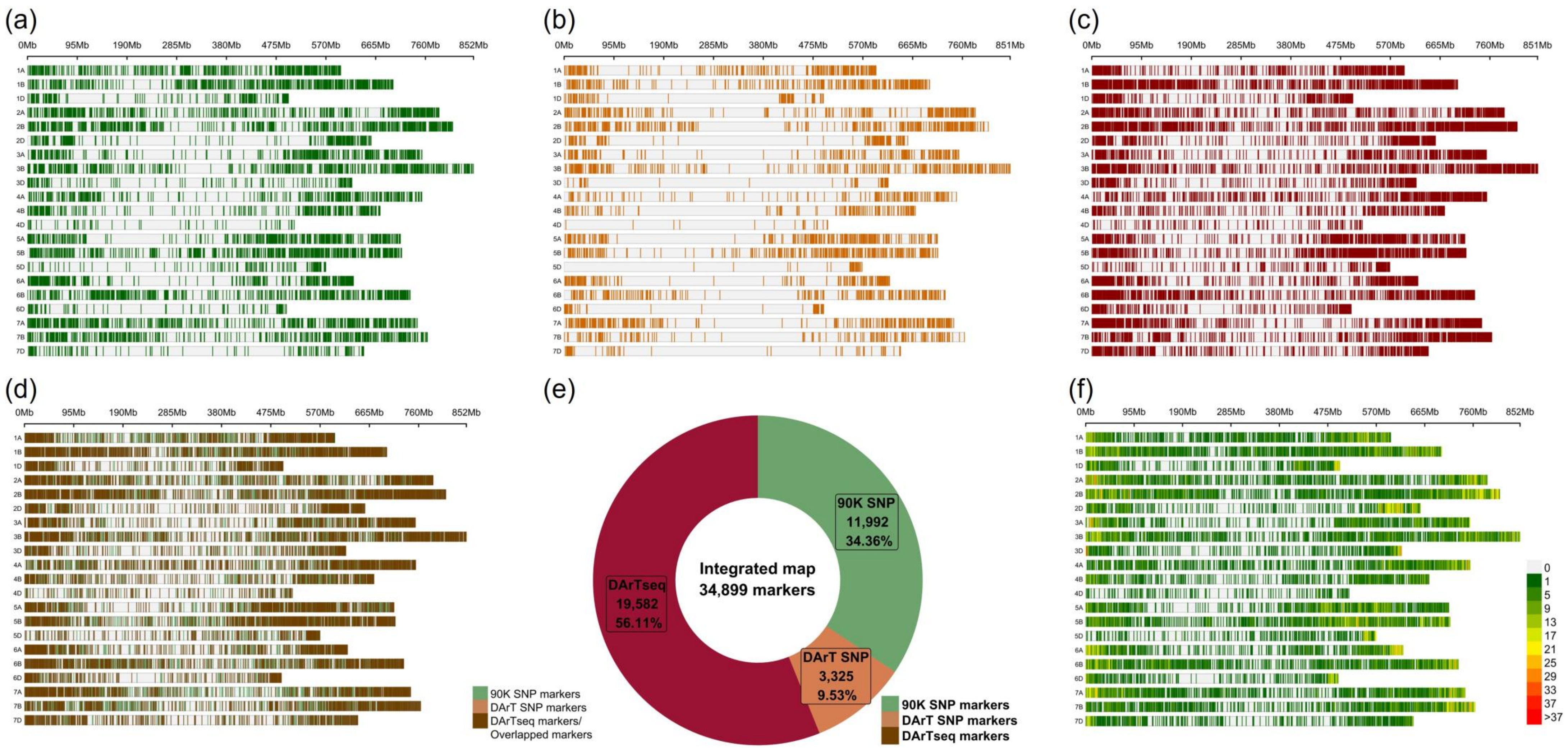


**Figure S1** Integrated marker map with 90K, DArT SNP, and DArTsilico (DArTseq) markers. **(a), (b),** and **(c)** show the distribution and density of 90K, DArT SNP, and DArTsilico (DArTseq) on chromosomes, respectively. **(d)** The integration of each type of markers on chromosomes. **(e)** The percentage of each type of markers to the integrated markers. **(f)** The distribution and density of integrated markers on chromosomes.

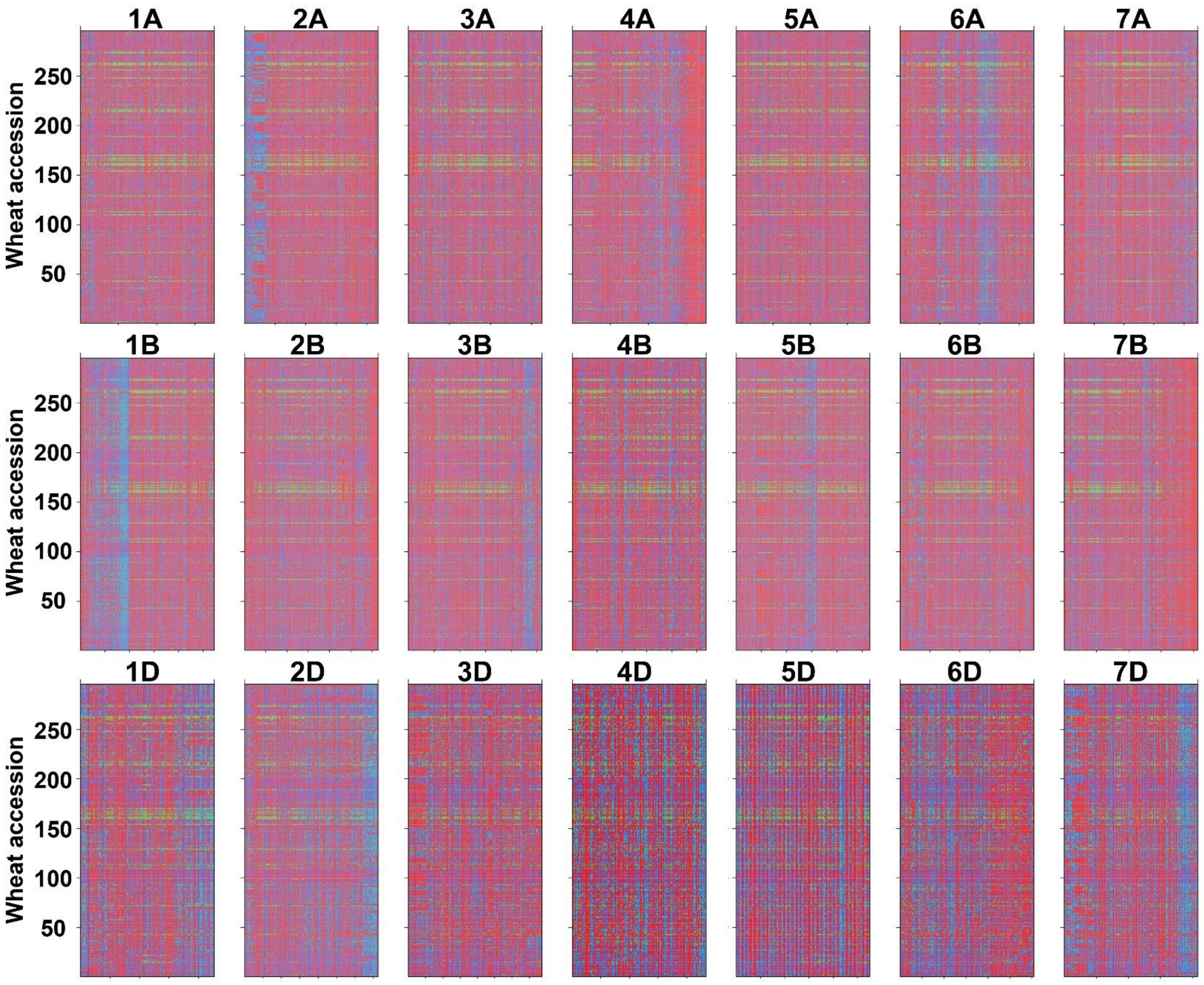


**Figure S2** Marker allele distributions of integrated map on each chromosome in 295 Vavilov wheat accessions. Red and blue represent two types of homozygous alleles, respectively. Green denotes missing/heterozygous genotype.

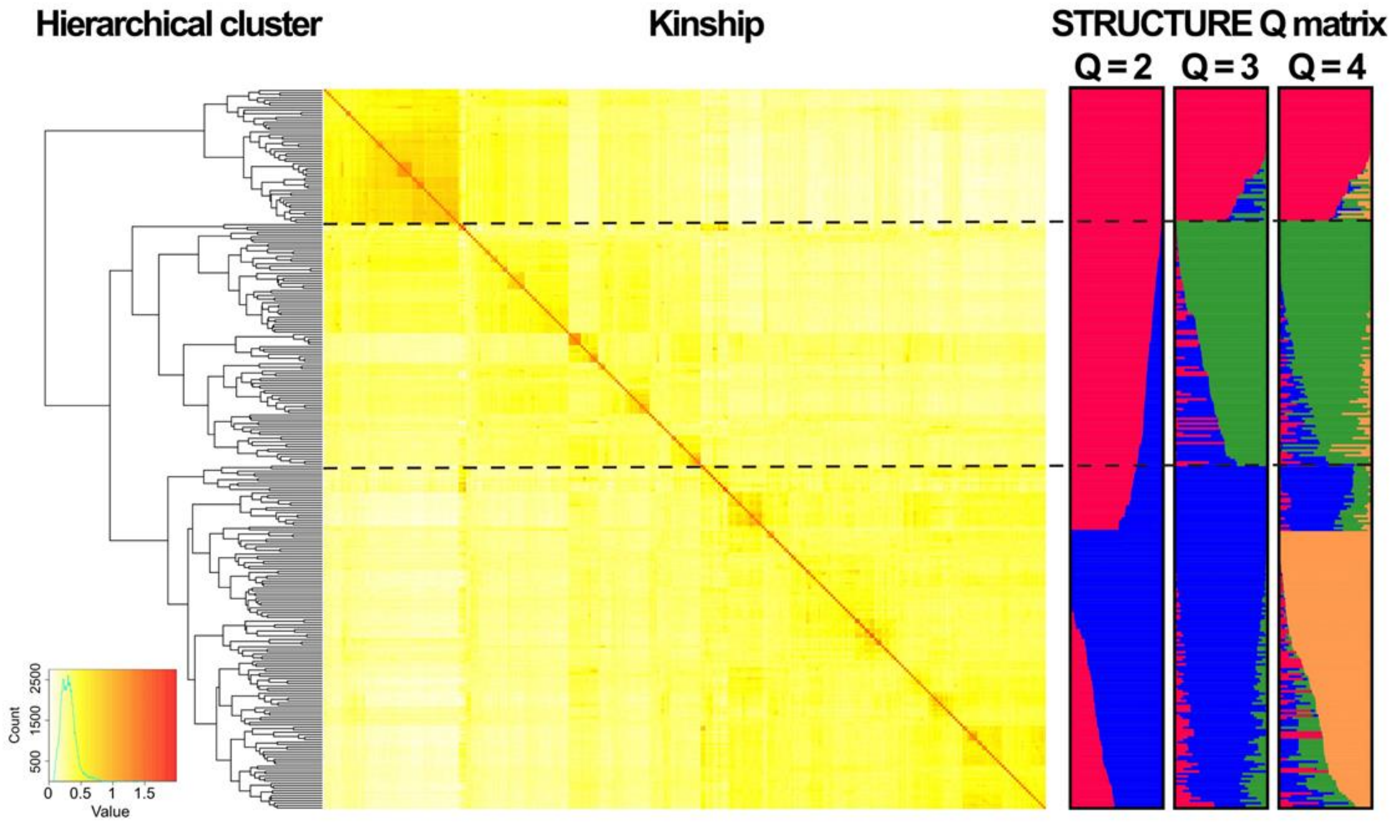


**Figure S3** Population structure of the 295 Vavilov wheat accessions revealed by Ward's hierarchical clustering, Kinship, and STRUCTURE software analyses. Dashed horizontal lines indicate genetic similarity thresholds used to classify the 295 wheat accessions into three main subgroups.

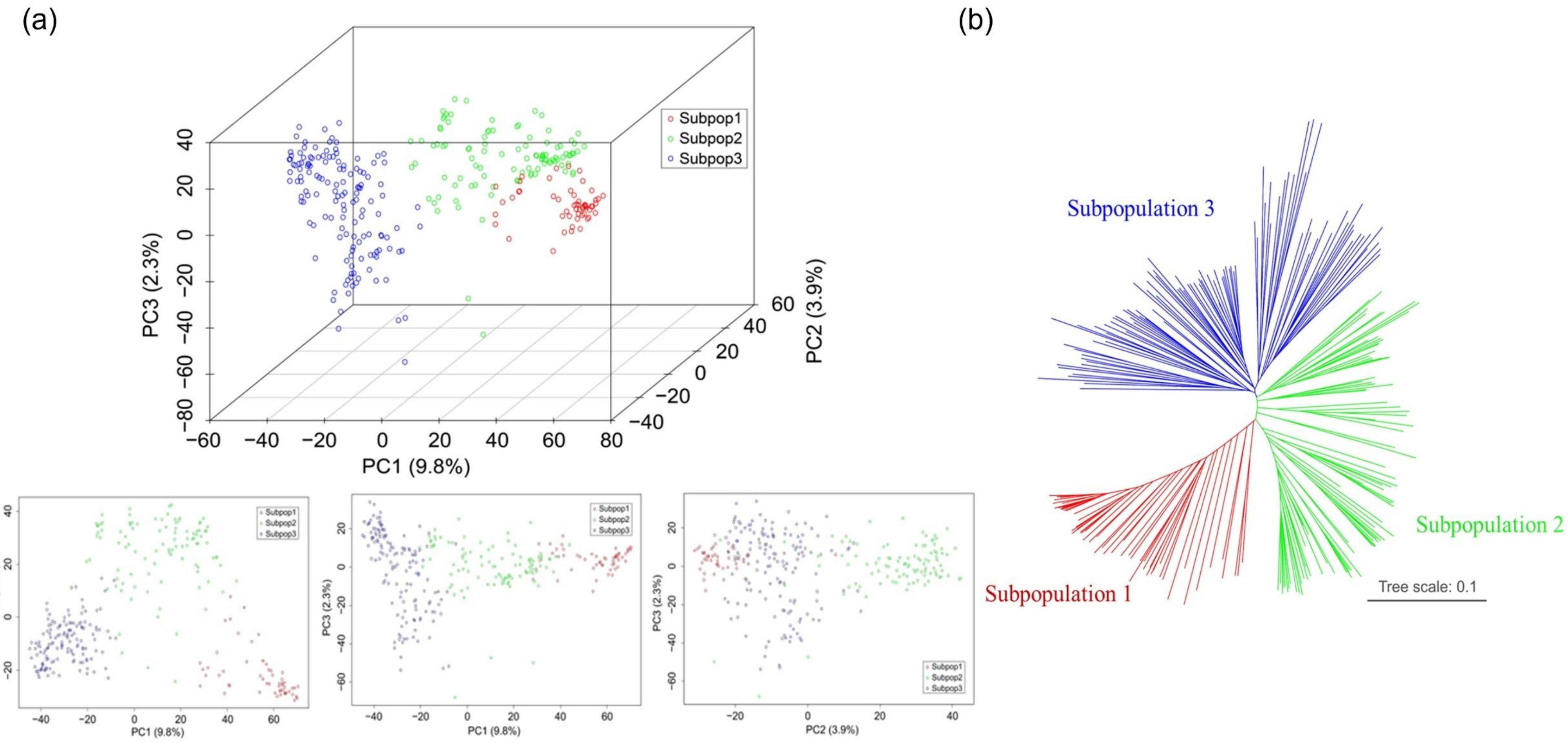


**Figure S4** Three subpopulations are classified in the Vavilov wheat panel. **(a)** Principal component analysis using the first three eigenvectors. **(b)** Neighbor-Joining tree analysis using genetic distance.

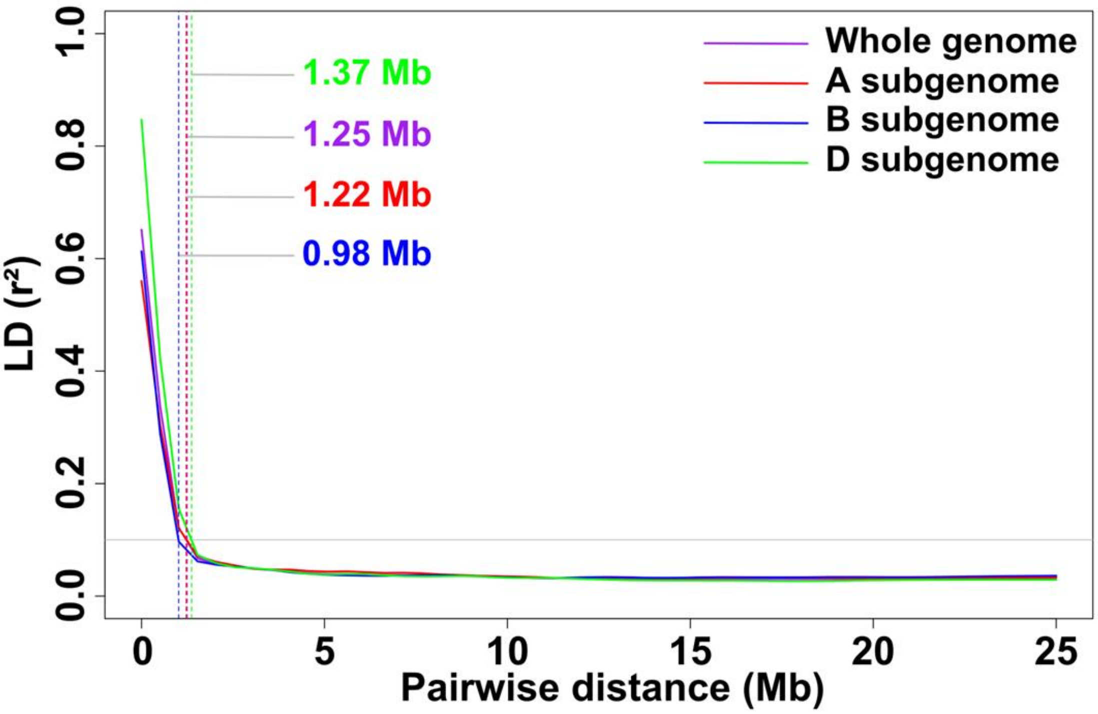


**Figure S5** Linkage disequilibrium (LD) decay over physical distances on the A, B, D subgenome, and the whole genome. Pairwise LD was estimated using the squared allele frequency correlation ($r^2$) of markers.

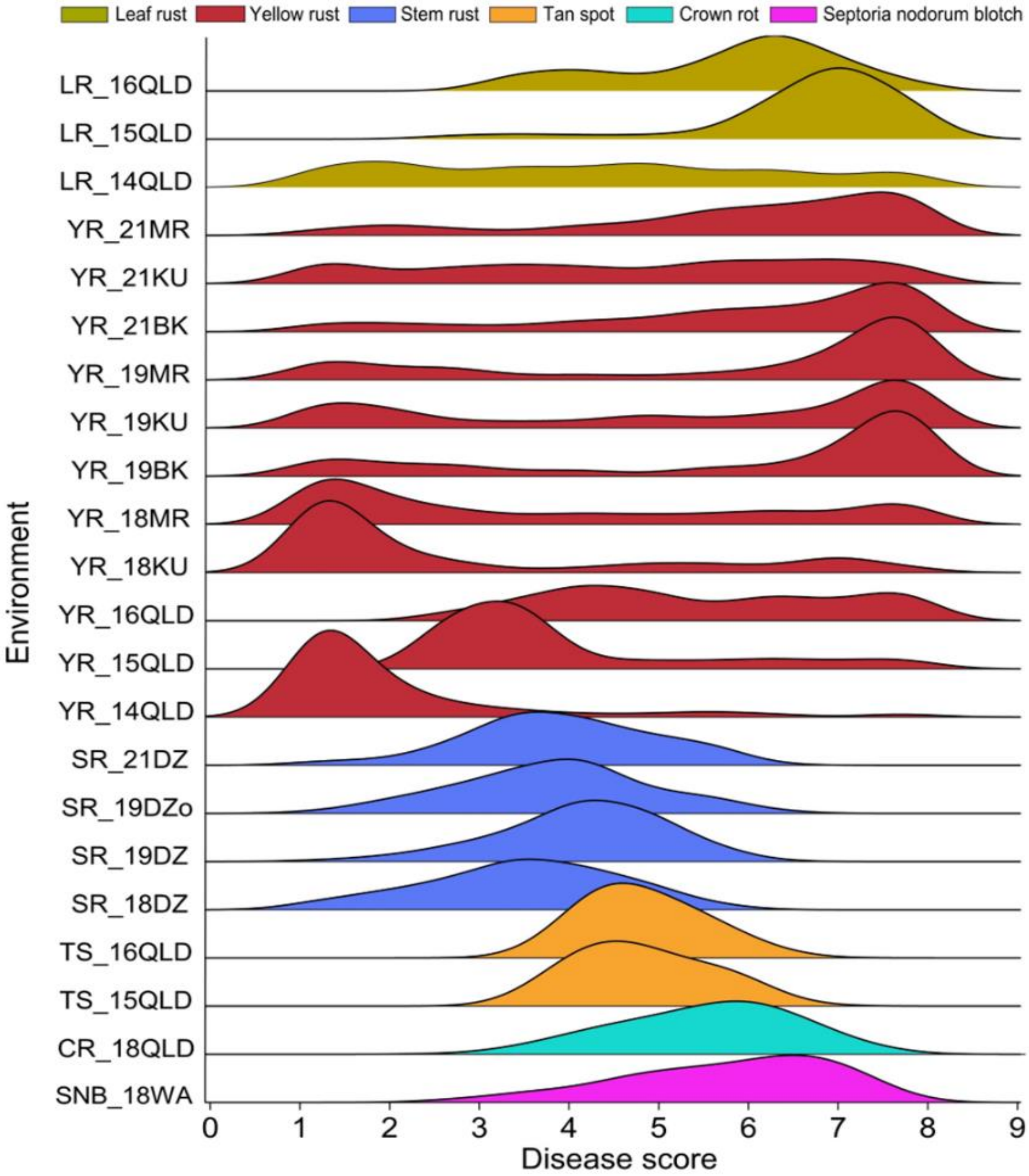


**Figure S6** Phenotypic variations and distributions of 22 individual experiments on disease responses to leaf rust (LR), yellow rust (YR), stem rust (SR), tan spot (TS), crown rot (CR) and Septoria nodorum blotch (SNB). Disease response data were converted to consistent scale 0-9 (0 = most resistant; 9 = most susceptible). Only the field dataset recorded in the last time-point was used in each environment.

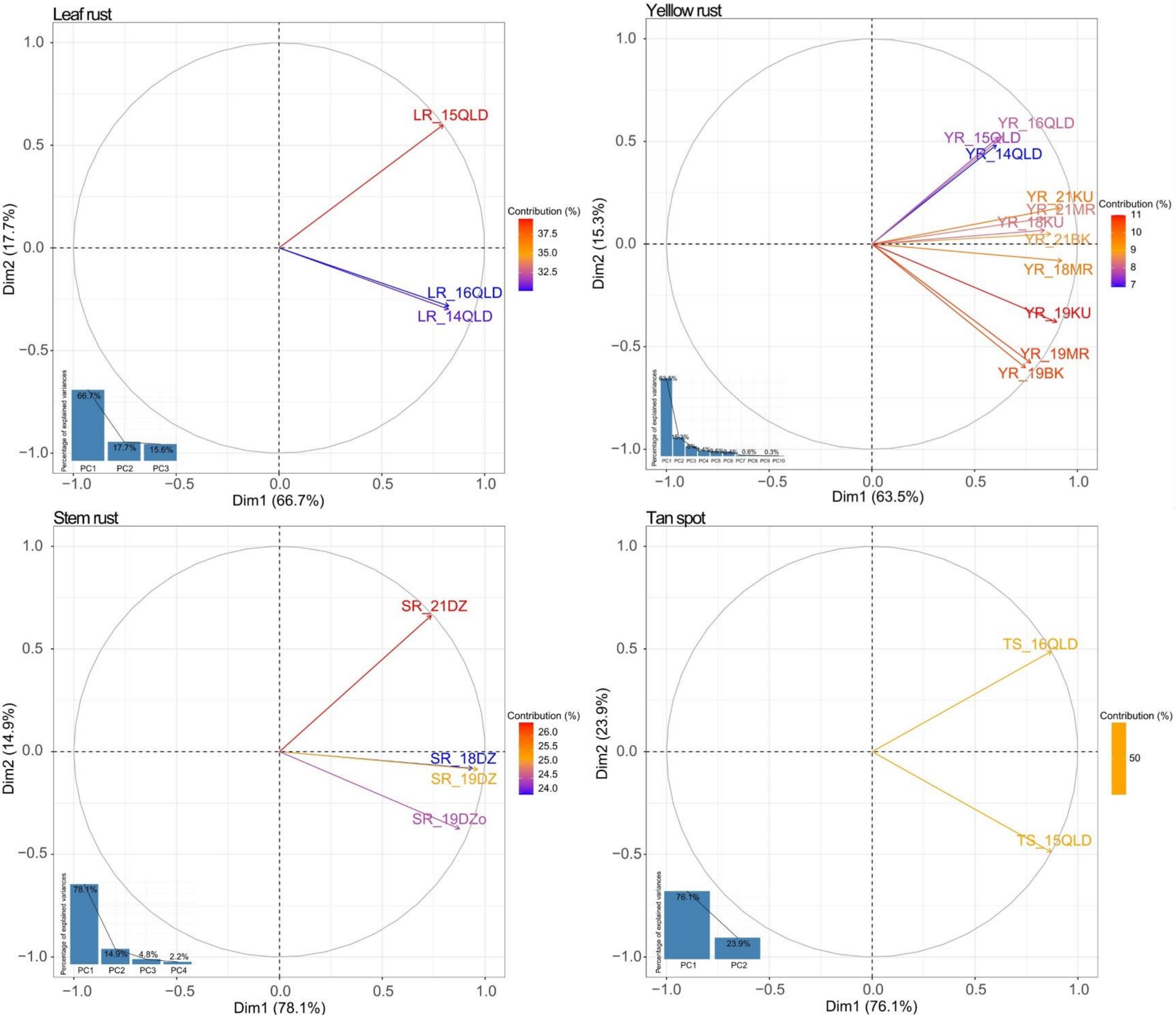


**Figure S7** Principal component analysis (PCA) of phenotypic datasets of leaf rust (LR), yellow rust (YR), stem rust (SR), and tan spot (TS).

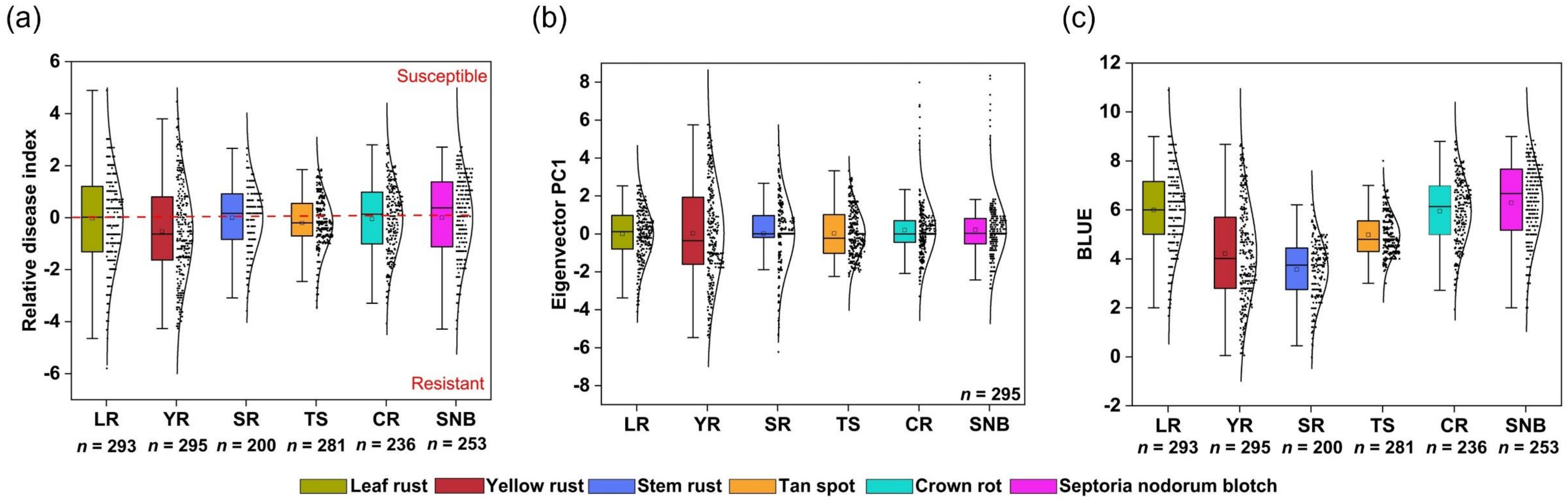


**Figure S8** Distributions of the three phenotypic parameters representing the resistance level of each wheat accession. **(a)** Relative disease index (Ri) values of different diseases. **(b)** Eigenvector principal component 1 (PC1) values of different diseases. Missing phenotypic values were imputed using expectation maximization (EM) method. **(c)** Best linear unbiased estimate (BLUE) values of different diseases. *n* represents the number of available phenotypic datapoints.

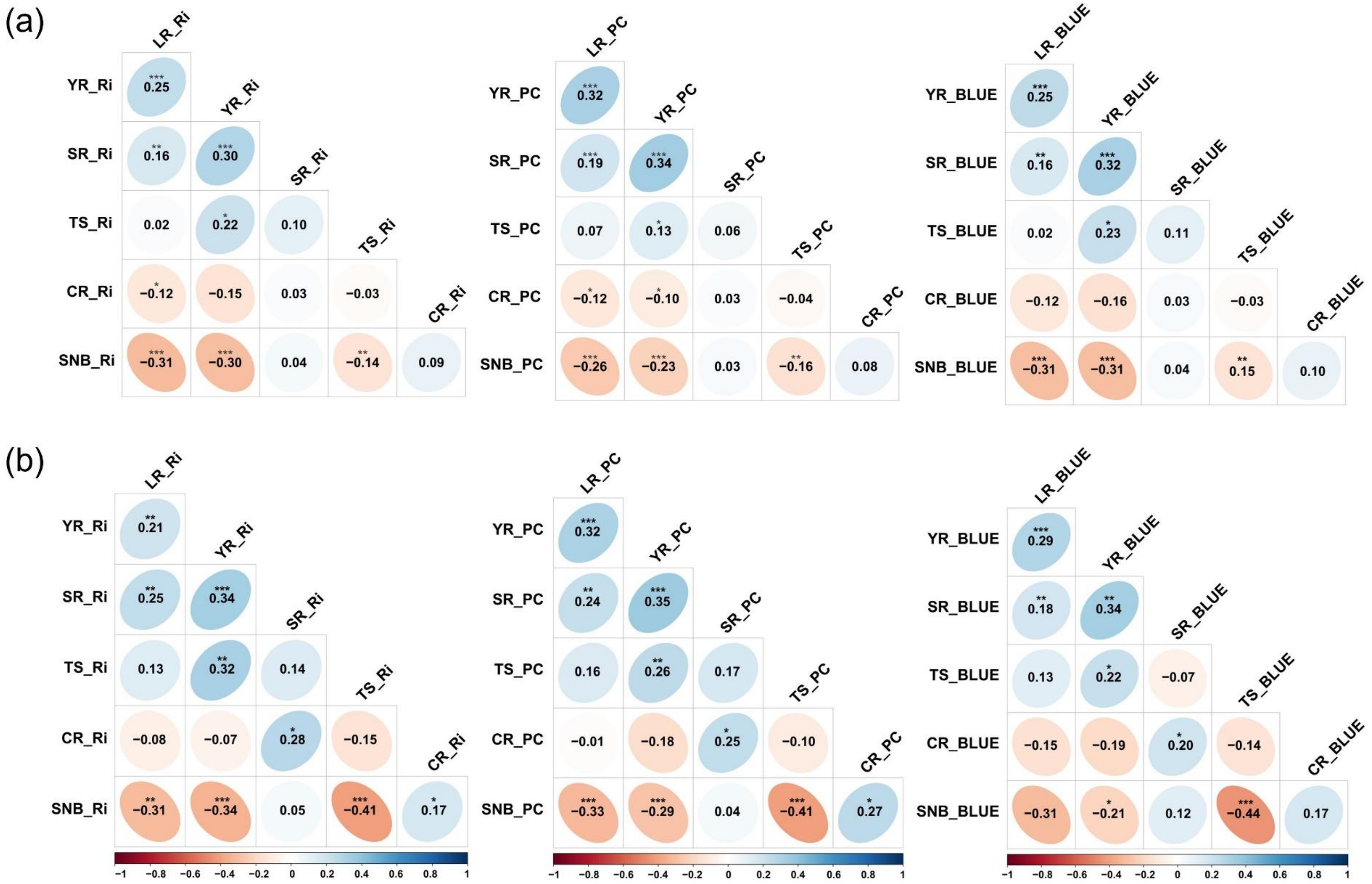


**Figure S9** Relationships between six wheat major diseases evaluated by three different phenotypic parameters, relative disease index (Ri), eigenvector principal component 1 (PC1), and best linear unbiased estimate (BLUE). **(a)** Pearson correlations between six wheat disease traits based on Ri, PC and BLUE, respectively. **(b)** Global genetic correlations between six wheat disease traits based on Ri, PC and BLUE, respectively. LR, leaf rust; YR, yellow rust; SR, stem rust; TS, tan spot; CR, crown rot; SNB, Septoria nodorum blotch. * $P < 0.1$, ** $P < 0.01$, *** $P < 0.001$.

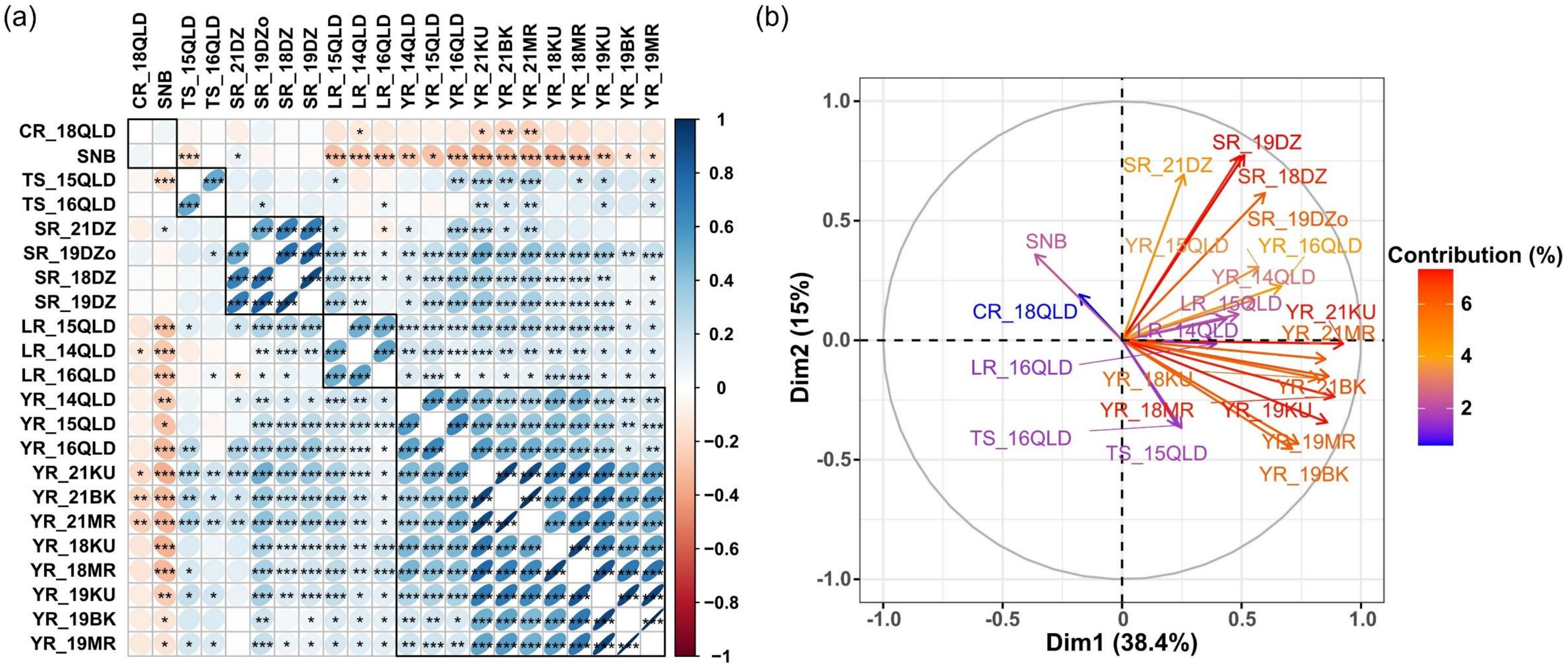


**Figure S10** Relationships between 22 individual experiments on disease responses to leaf rust (LR), yellow rust (YR), stem rust (SR), tan spot (TS), crown rot (CR) and Septoria nodorum blotch (SNB). **(a)** Pearson correlations with Ward's hierarchical clustering. * $P < 0.1$, ** $P < 0.01$, *** $P < 0.001$. **(b)** Two-dimension principal component analysis.

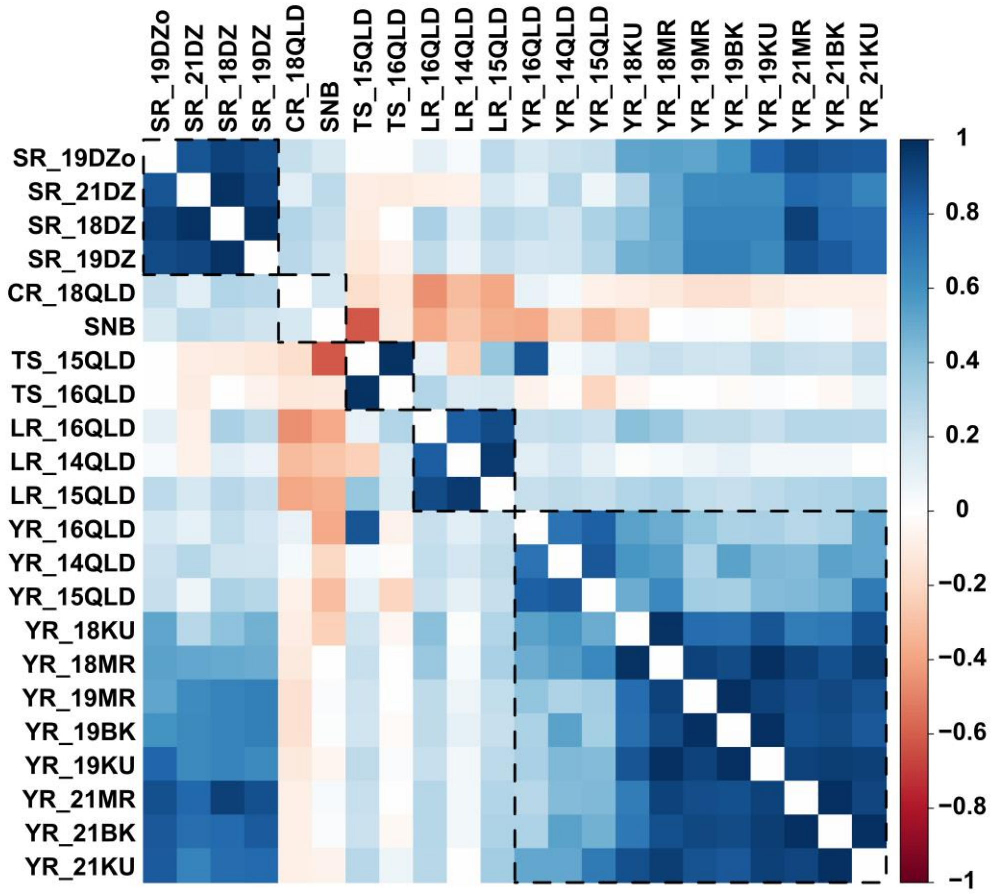


**Figure S11** Genetic correlations among 22 individual experiments on disease responses to leaf rust (LR), yellow rust (YR), stem rust (SR), tan spot (TS), crown rot (CR) and Septoria nodorum blotch (SNB).

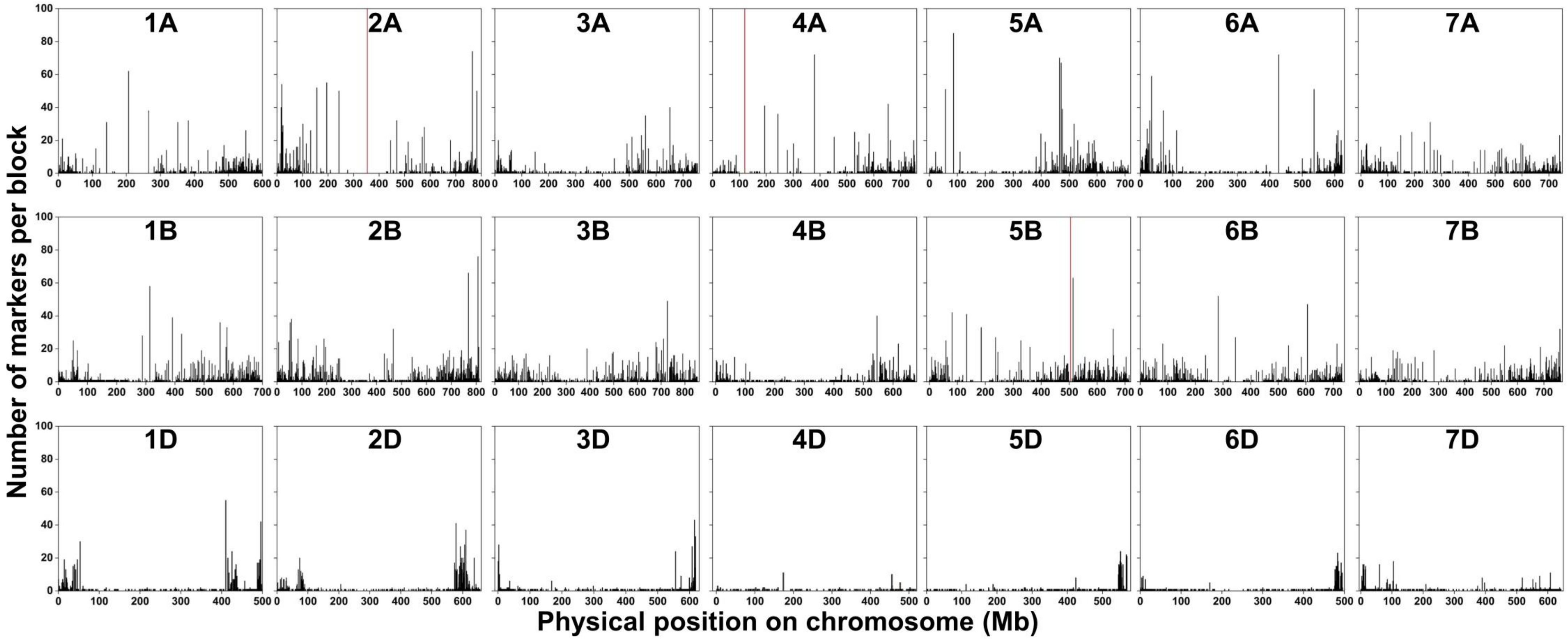


**Figure S12** Genome-wide distribution and size of defined haploblocks. Red bars on 2A, 4A, and 5B indicate a large linkage disequilibrium (LD) peak comprising 151, 123, and 102 markers, respectively, which exceeds the y-axis scale. Physical positions and sizes of haploblocks were defined according to the positions of included markers on IWGSC Refseqv2.1 and the number of markers per block, respectively.

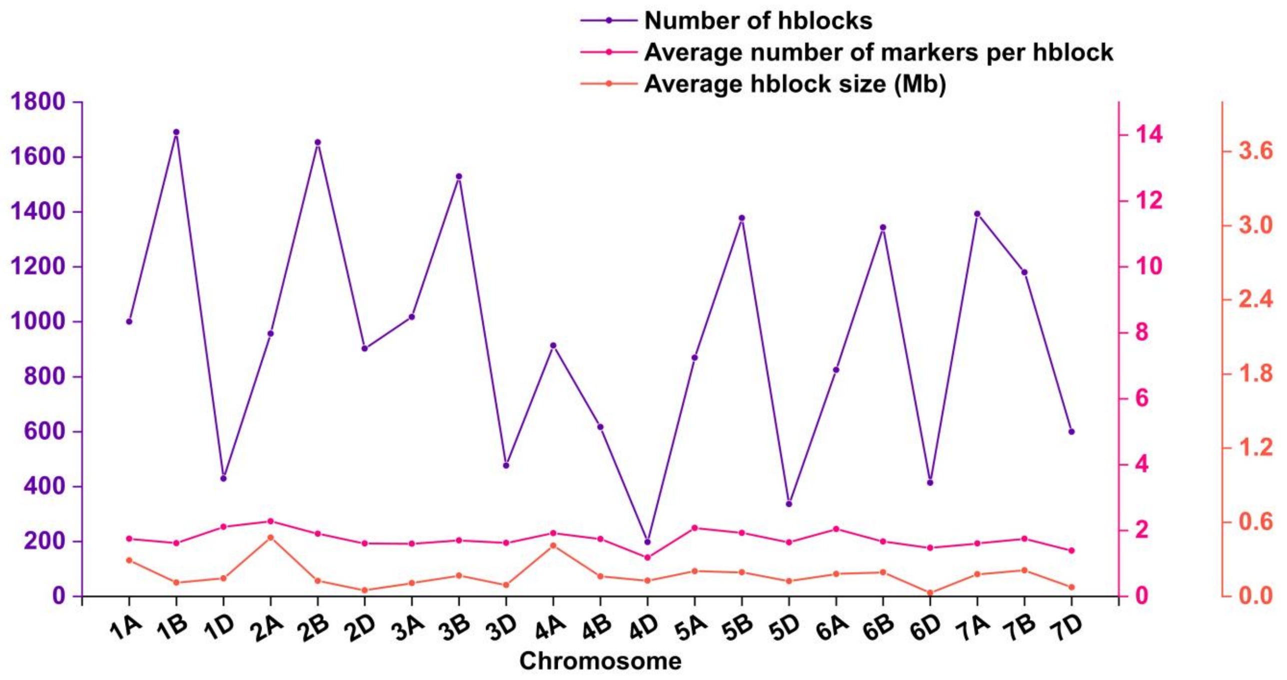


**Figure S13** Distribution patterns of haploblocks across all the 21 wheat chromosomes. The total number of blocks, average number of markers per block, and average block size in mega base (Mb) are showed on each of the wheat chromosomes.

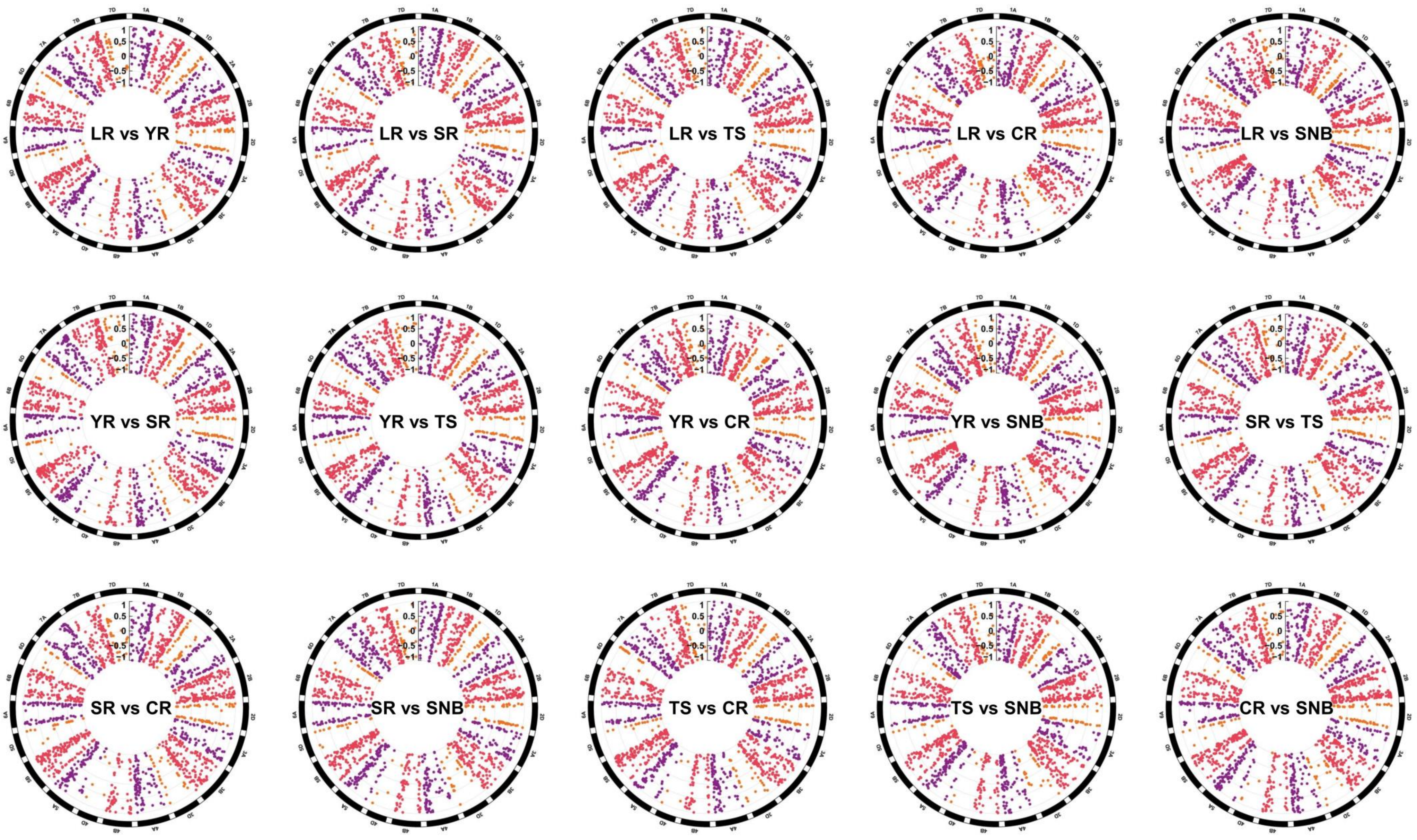


**Figure S14** Genome-wide plot of the local genetic correlation estimates for 15 disease trait pairs. Genome-wide 1,756 hblocks harboring >3 markers were used to calculate correlations (*r*) between all 15 pairs of disease traits. Only *r* estimates with Bonferroni-corrected *P*-value <0.05 were considered as significant. LR, leaf rust; YR, yellow rust; SR, stem rust; TS, tan spot; CR, crown rot; SNB, Septoria nodorum blotch.

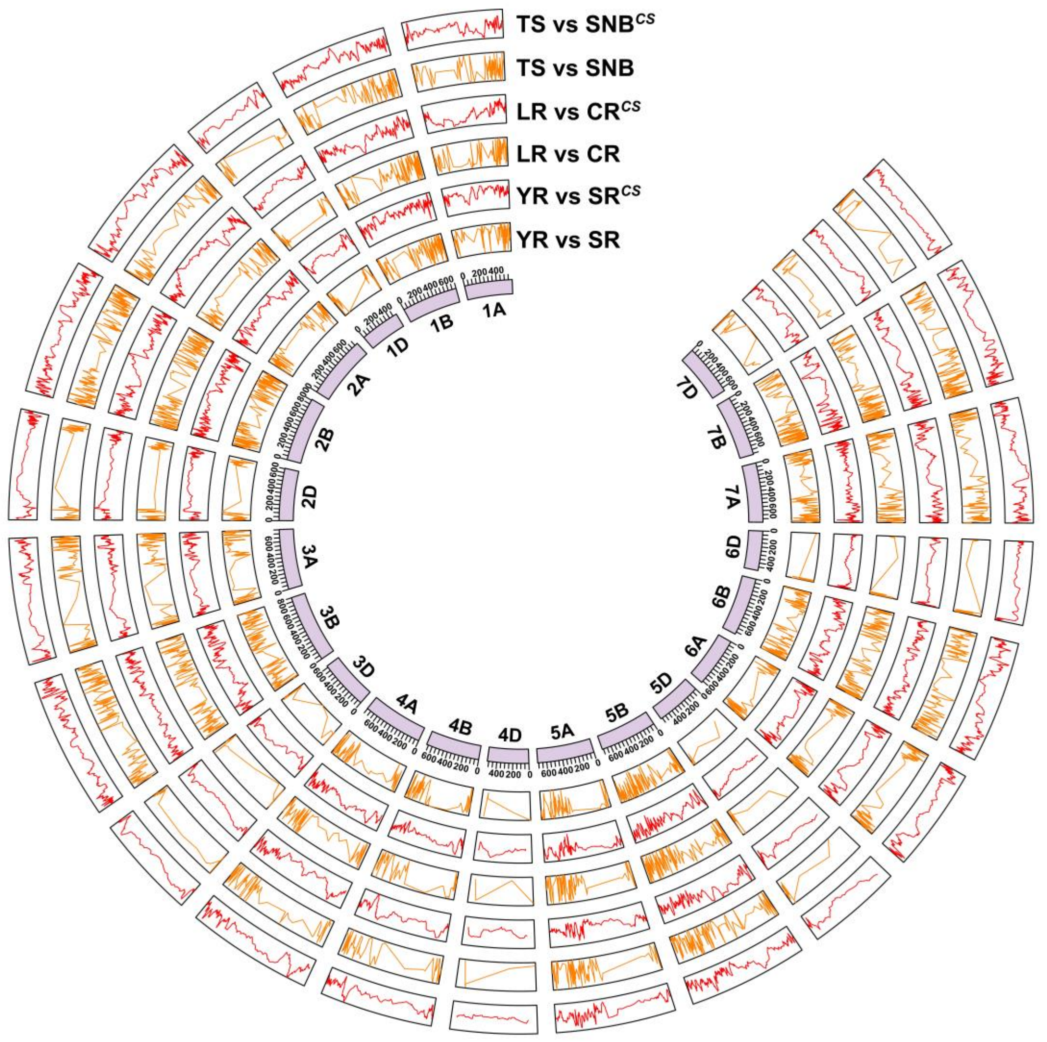


**Figure S15** Comparison of local genetic correlation calculations between haploblock-based method and correlation scan (CS). CS used correlations of effects of 500-marker sliding windows with a step length of 100 markers to identify local genomic regions that either drive or antagonize the genetic correlations between traits. Three trait pairs with positive (YR vs SR), neutral (LR vs CR) and negative genetic correlations (TS vs SNB) were used to compare the results from haploblock-based method and marker-sliding-window-based CS. Similar genome-wide patterns are observed on most chromosomes for local genetic correlations from the two methods. Y axis represents correlation coefficient. Red, marker-sliding-window-based correlations. Orange, hblock-based correlations. LR, leaf rust; YR, yellow rust; SR, stem rust; TS, tan spot; CR, crown rot; SNB, Septoria nodorum blotch.

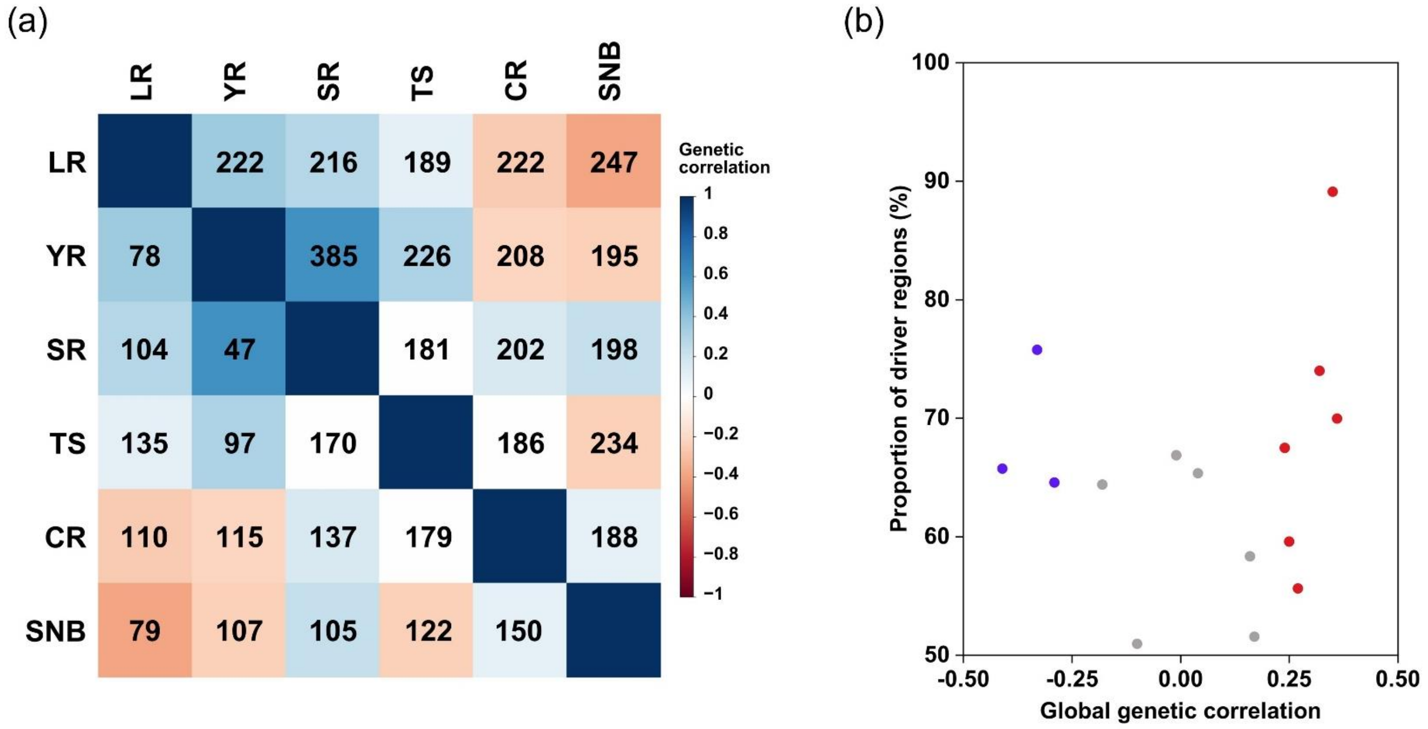


**Figure S16** Significant driving and antagonizing haploblocks along with genetic correlations between all 15 trait pairs. **(a)** Heatmap illustrating the number of significant driving (top) and antagonizing (bottom) haploblocks. The driver regions refer to the blocks driving the global genetic correlation, that is, the positive/ negative correlations of significant correlated blocks have the same direction with positive/ negative global genetic correlations of trait pairs, while the antagonizing regions have the opposite direction. The colours in the heatmap reflect the mean local correlations across all significant haploblocks. LR, leaf rust; YR, yellow rust; SR, stem rust; TS, tan spot; CR, crown rot; SNB, Septoria nodorum blotch. **(b)** Dot plot showing the proportion of driver regions along with global genetic correlations between all 15 trait pairs. The red points denote the trait pairs that have significant positive genetic correlations, while the blue represent significant negative genetic correlations.

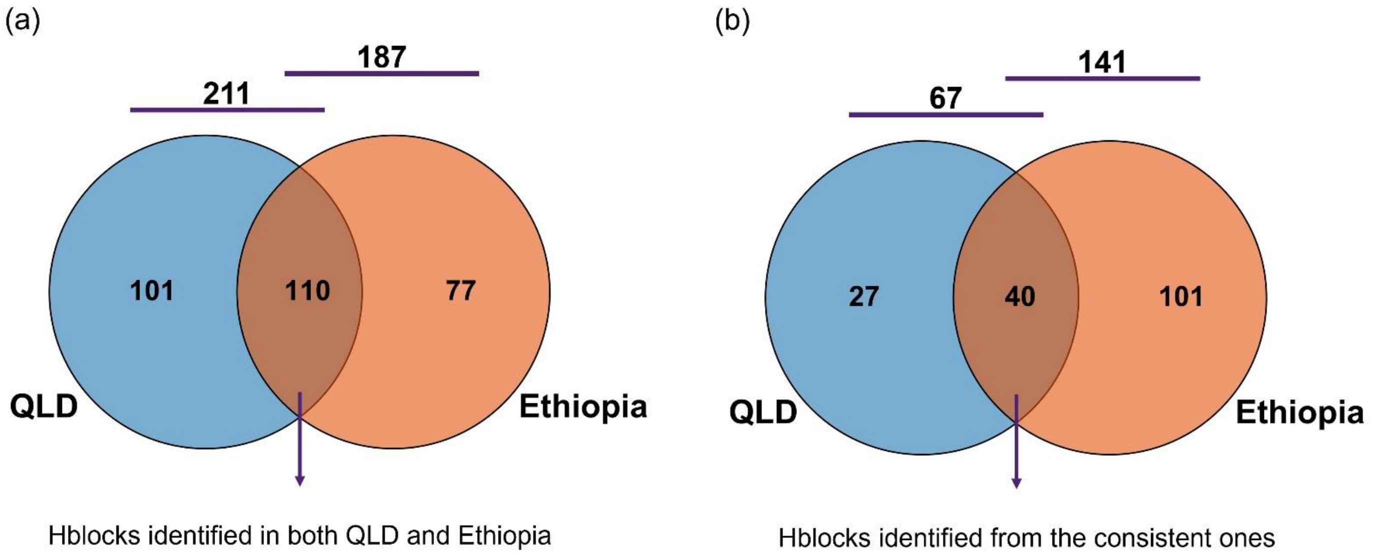


**Figure S17** Haploblocks contribute to high yellow rust (YR) correlations within and between Queensland (QLD) and Ethiopia environments. **(a)** Venn plot showing the total number of haploblocks relevant to YR in QLD and Ethiopia environments. **(b)** Venn plot showing the number of consistent YR haploblocks identified in QLD and Ethiopia environments. The blocks identified at least two environments in each QLD and Ethiopia were regarded as the consistent.

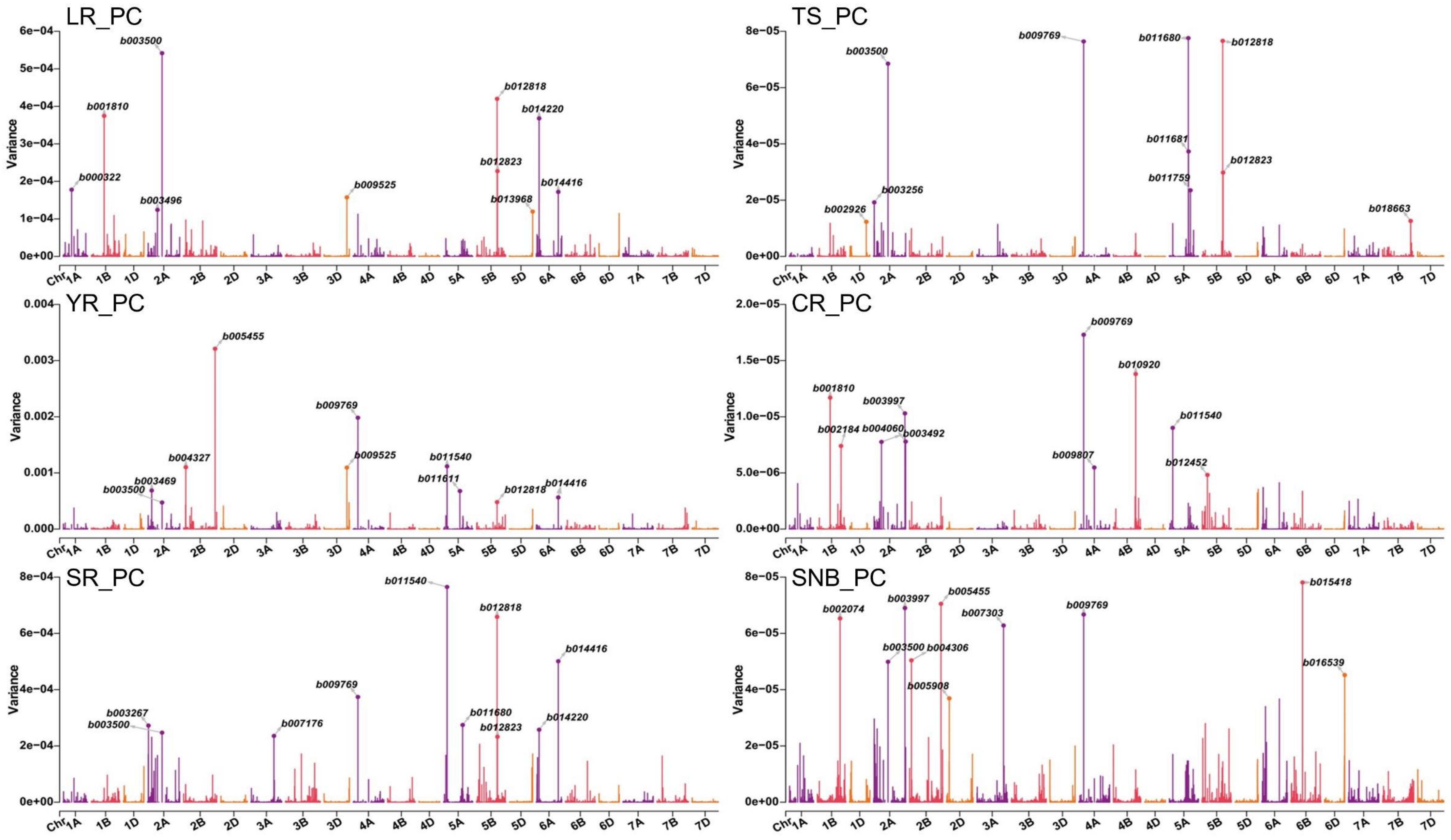

**Figure S18** Whole-genome haploblock variances for the six diseases using eigenvector principal component 1 (PC1). Top 10 haploblocks with the highest variances are marked for each respective disease. LR, leaf rust; YR, yellow rust; SR, stem rust; TS, tan spot; CR, crown rot; SNB, Septoria nodorum blotch.

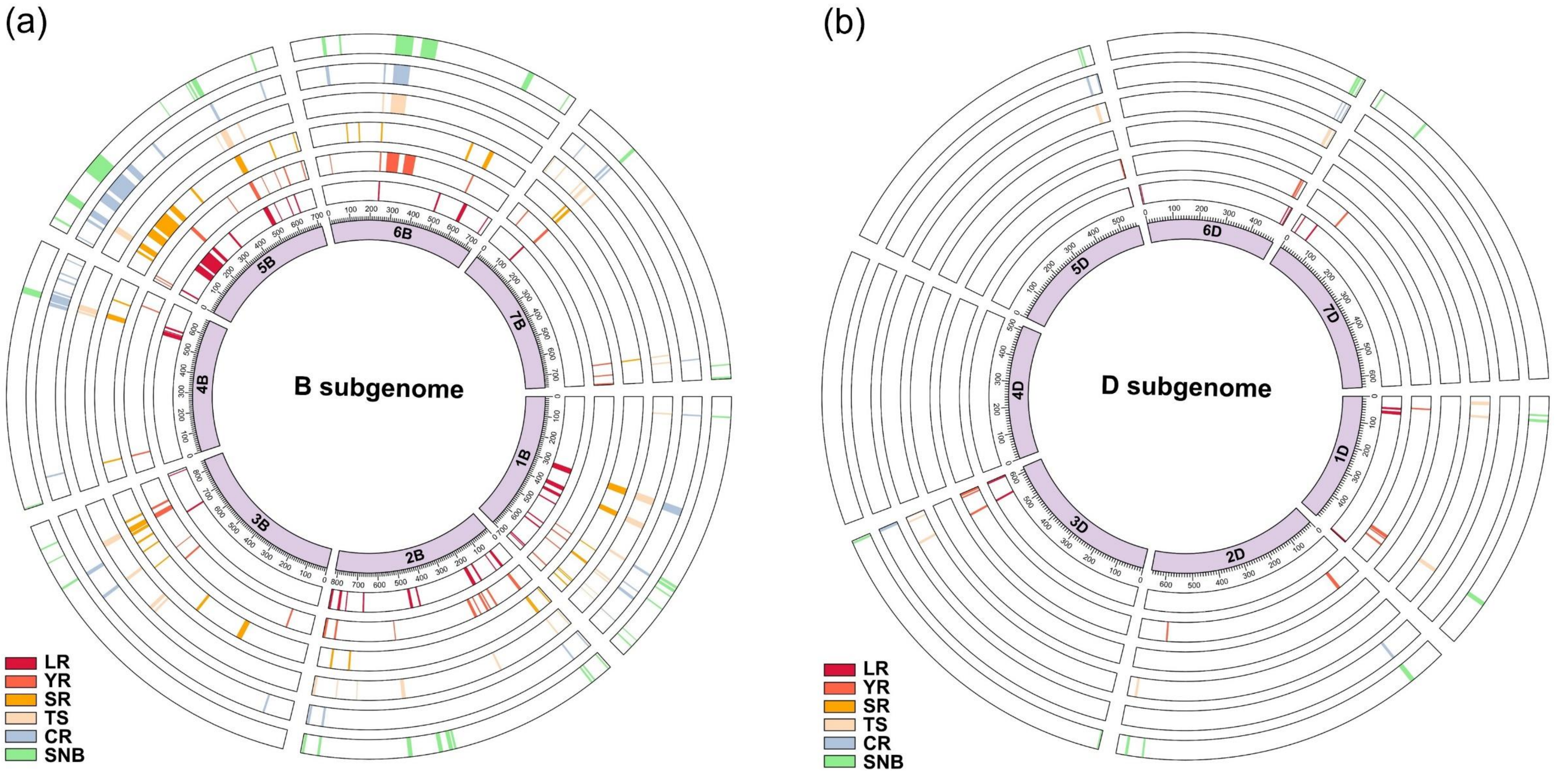


**Figure S19** Circos plot showing the chromosome distributions of the top 100 haploblocks on B and D subgenomes for each of the six diseases. **(a)** B subgenome distributions of top 100 blocks with the highest haplotype variances for each respective disease. **(b)** D subgenome distributions of top 100 blocks with the highest haplotype variances for each respective disease. LR, leaf rust; YR, yellow rust; SR, stem rust; TS, tan spot; CR, crown rot; SNB, Septoria nodorum blotch.

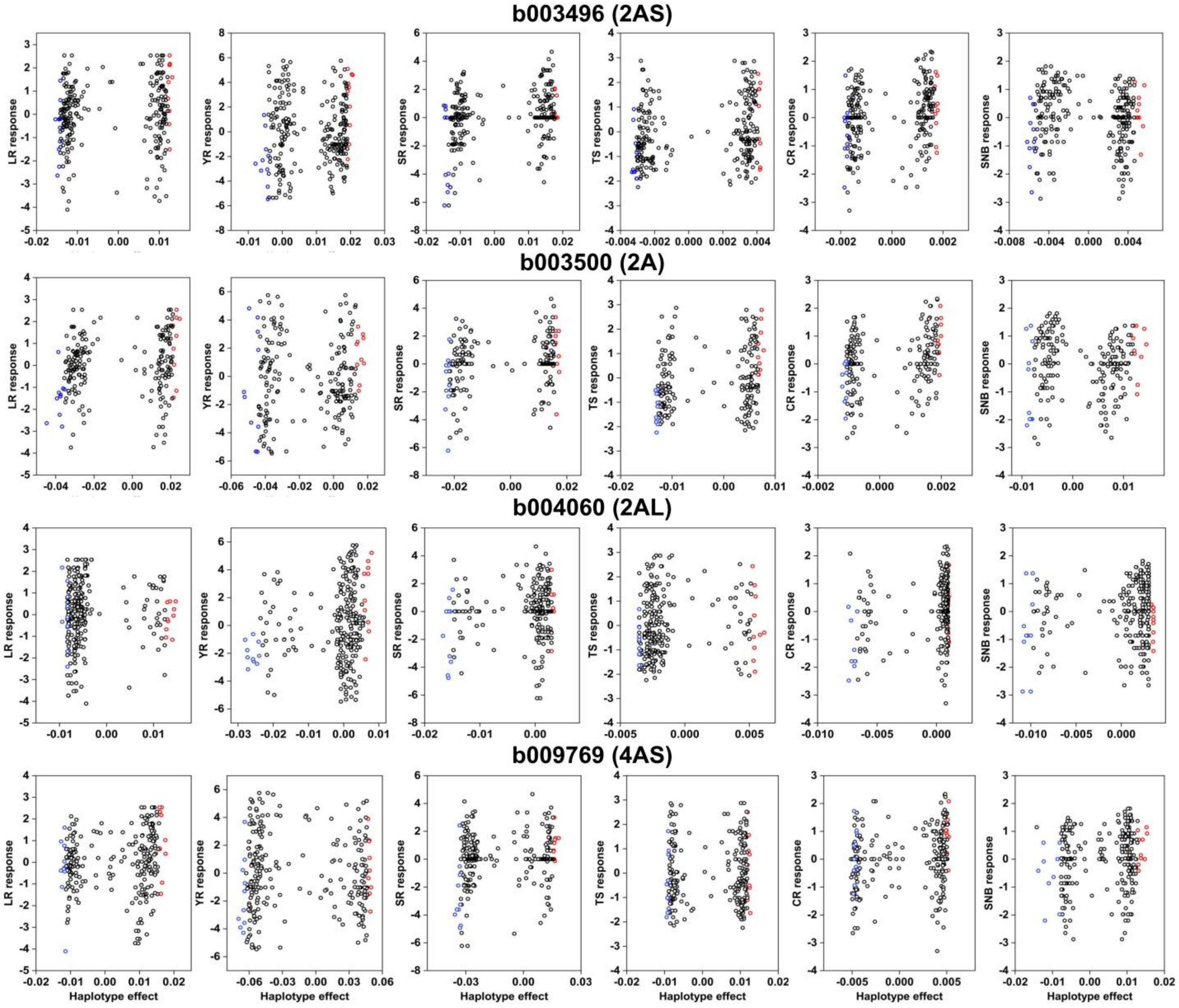


**Figure S20** Phenotypic effects contributed by haplotypes of the four haploblocks, b003496, b003500, b004060 and b009769 related to six diseases. Each point represents one wheat accession from the Vavilov panel. Red and blue points represent ten wheat accessions with the highest and lowest haplotype effects of each hblock for each disease, respectively. Y axis is disease response represented by eigenvector principal component 1 (PC1) of each trait. Marked phenotypic difference are observed between most red and blue pairs. LR, leaf rust; YR, yellow rust; SR, stem rust; TS, tan spot; CR, crown rot; SNB, Septoria nodorum blotch.

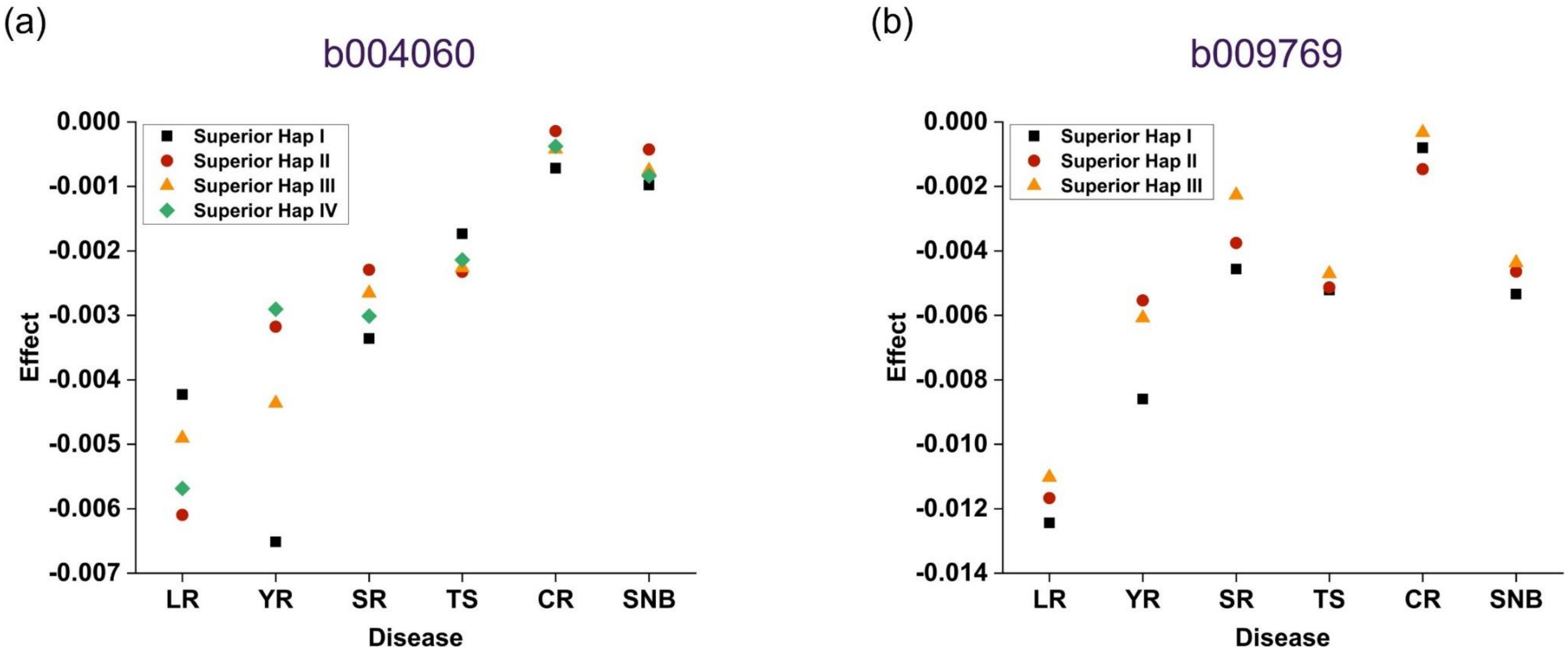


**Figure S21** Superior haplotypes of **(a)** b004060 and **(b)** b009769 with positive effects to improve resistance levels of the six diseases. LR, leaf rust; YR, yellow rust; SR, stem rust; TS, tan spot; CR, crown rot; SNB, Septoria nodorum blotch.

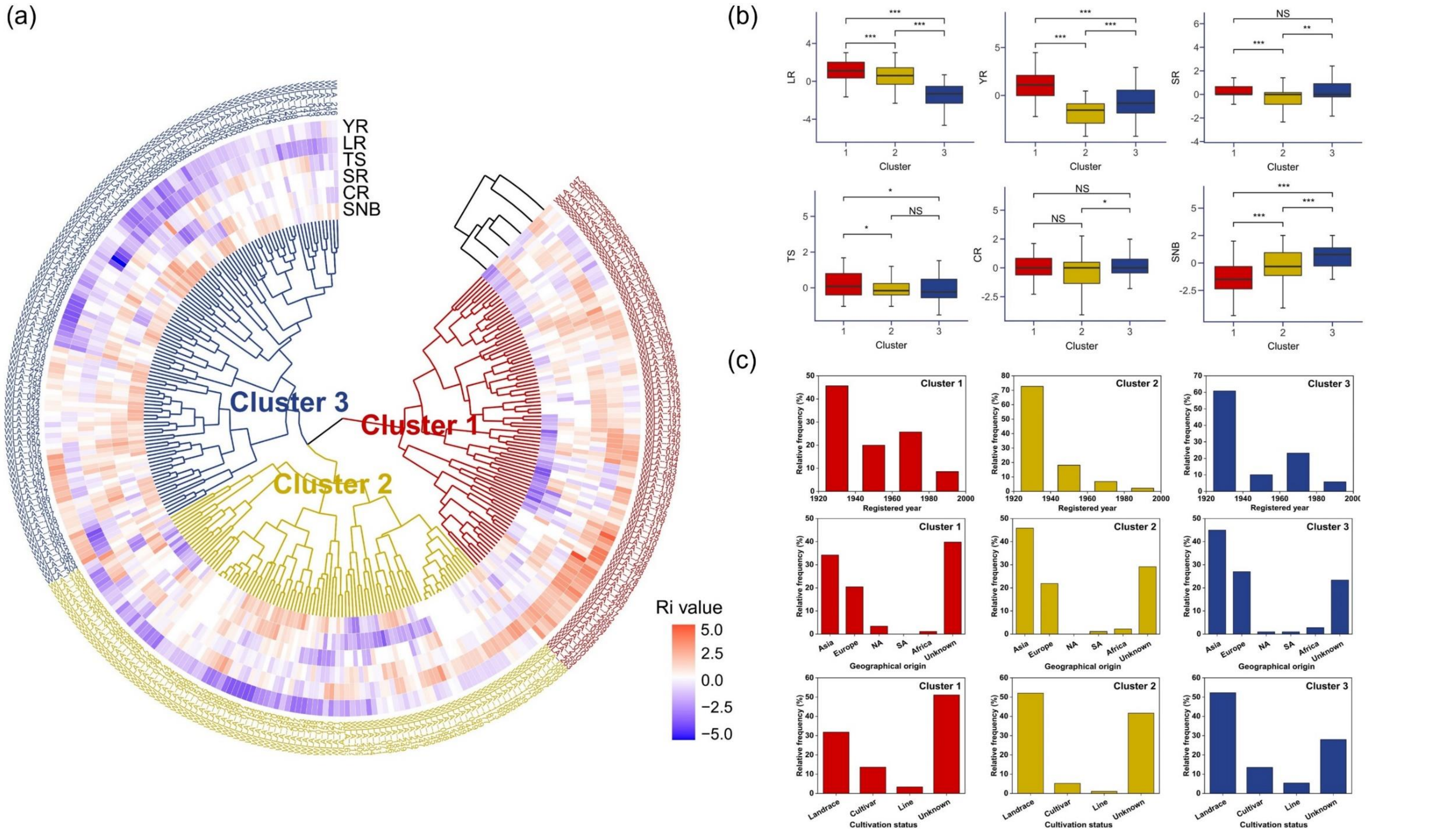


**Figure S22** Screening of Vavilov wheat accessions based on relative disease index (Ri) of six diseases. **(a)** Clustering heatmap of diverse wheat accessions based on six diseases. LR, leaf rust; YR, yellow rust; SR, stem rust; TS, tan spot; CR, crown rot; SNB, Septoria nodorum blotch. **(b)** Comparison of classified clusters for each respective disease. **(c)** Registered year, geographical origin and cultivation status of the different clusters. NA, North America; SA, South America.

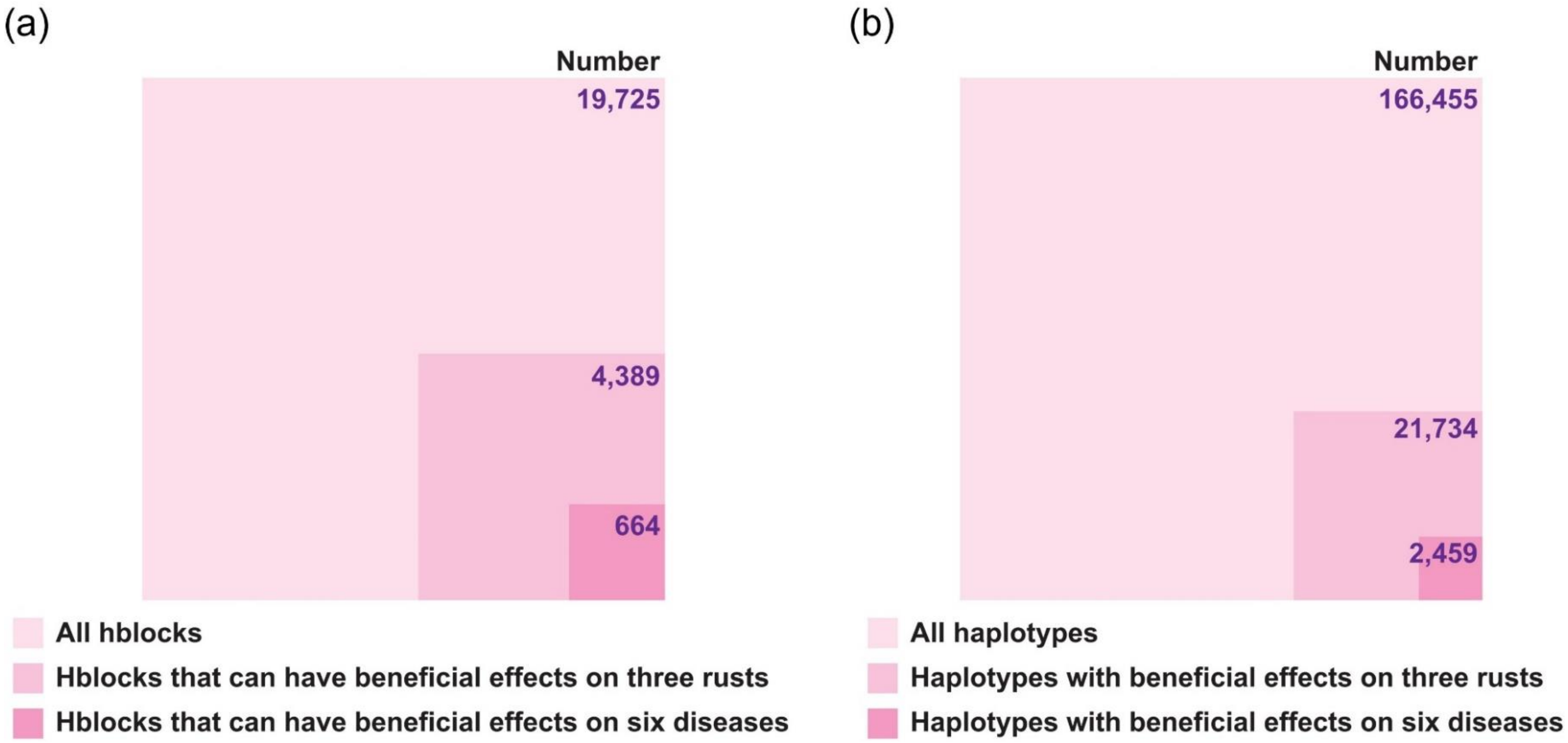


**Figure S23** Number of **(a)** haploblocks and **(b)** corresponding haplotypes that can have beneficial effects on three rusts and six diseases. All of 19,725 haploblocks and corresponding 166,455 haplotypes with minor or major effects were analysed.

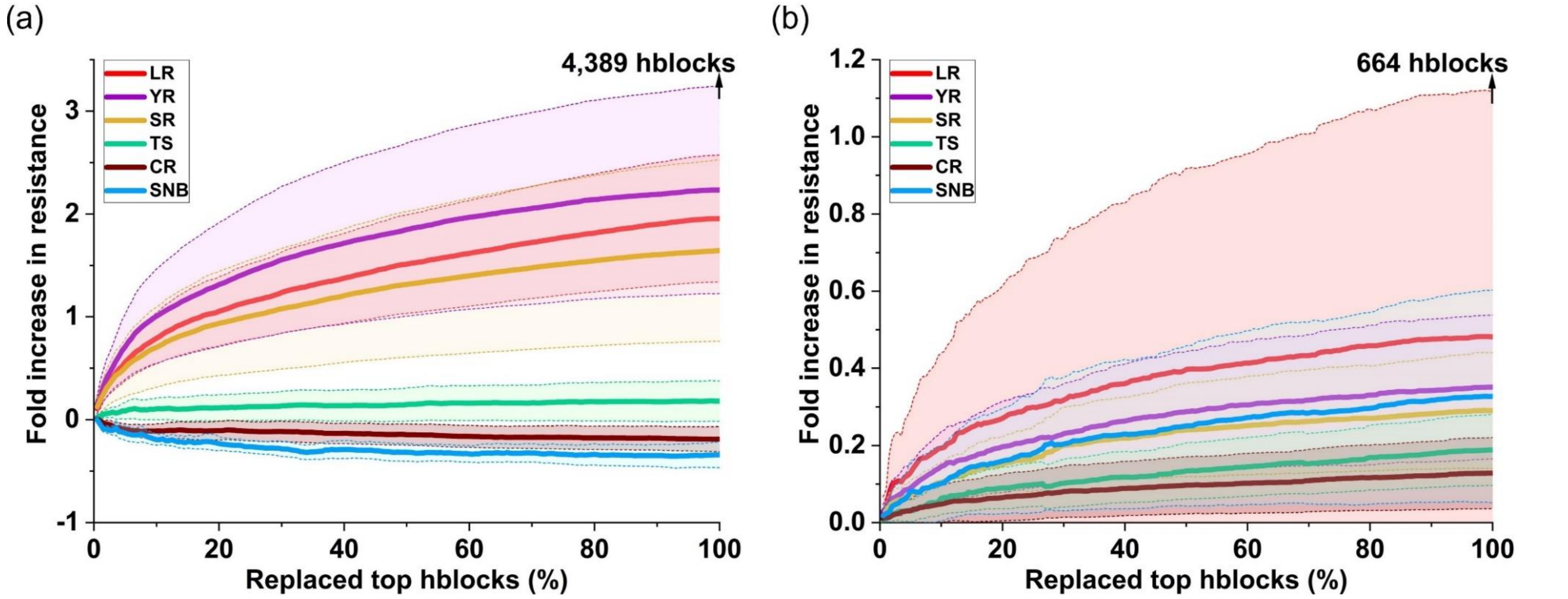


**Figure S24** Resistance predictions evaluating effects of stacking the most beneficial haplotypes to **(a)** three rusts and **(b)** six diseases on multiple disease resistance (MDR). 4,389 three-rust and 664 six-disease haploblocks were selected for the haplotype-stacking simulation, respectively. Nineteen out of 295 wheat accessions screened with high resistance to at least five diseases were used as a basis. *In silico* genotypes were generated from these selected cultivars by exchanging the original haplotypes by the most favourable haplotypes at the hblocks. Genomic estimated breeding values (GEBV) relative changes were calculated. The solid lines indicate the mean of predicted changes, and standard errors are showed by colored shadows with dashed lines. LR, leaf rust; YR, yellow rust; SR, stem rust; TS, tan spot; CR, crown rot; SNB, Septoria nodorum blotch.
